# Structural characterization of antibody-responses from Zolgensma treatment provides the blueprint for the engineering of an AAV capsid suitable for redosing

**DOI:** 10.1101/2024.05.01.590489

**Authors:** Mario Mietzsch, Austin R. Nelson, Jane Hsi, Jon Zachary, Lindsay Potts, Paul Chipman, Mohammad Ghanem, Neeta Khandekar, Ian E. Alexander, Grant J. Logan, Juha T. Huiskonen, Robert McKenna

## Abstract

Monoclonal antibodies (mAbs) are useful tools to dissect the neutralizing antibody response against the adeno-associated virus (AAV) capsids used as gene therapy delivery vectors. This study structurally characterizes the interactions of 21 human-derived antibodies from patients treated with the AAV9 vector, Zolgensma^®^, utilizing high-resolution cryo-electron microscopy. The majority of the bound antibodies do not conform to the icosahedral symmetry of the capsid, thus requiring localized reconstructions. These complex structures provide unprecedented details of the mAbs binding interfaces, with some antibodies inducing structural perturbations of the capsid upon binding. Key surface capsid amino acid residues were identified facilitating the design of capsid variants with an antibody escape phenotype, with the potential to expand the patient cohort treatable with AAV9 vectors to include those that were previously excluded due to their pre-existing neutralizing antibodies, and possibly also to those requiring redosing.

## Introduction

Adeno-associated viruses (AAVs) have become the leading therapeutic gene delivery vector, as they are non-pathogenic, possess the ability to package and express therapeutic genes in a wide range of cell and tissue types^1-3^. They can mediate long-term correction of monogenic diseases and have been shown to be effective in numerous clinical trials. These successes have resulted in the approval of currently seven AAV-based biologics, including Zolgensma, an AAV9-based therapy for the treatment of spinal muscular atrophy^4-7^. The AAV vectors are composed of non-enveloped T=1 icosahedral capsids, consisting of 60 copies of the viral proteins (VP), and package single-stranded DNA transgene cassettes^8^.

The structures of several AAV capsids have been determined by X-ray crystallography and/or cryo-electron microscopy (cryo-EM)^8-10^. All AAV capsids VPs, consist of a conserved jelly-roll core and variable surface loops/regions (VR)^8^. The capsids are assembled via 2-, 3-, and 5-fold symmetry-related VP interactions^11^. The icosahedral 5-fold axes are surrounded by five loops that form the cylindrical channels which connect the interior to the exterior environment. Another characteristic feature of the capsids are protrusions surrounding the 3-fold axes. At the 2-fold axes and surrounding the 5-fold channels, the AAV capsids possess depressions that are separated by raised regions termed 2/5-fold walls. Structural variations between different AAV capsids arise from differences in amino acid (aa) sequences resulting in alternative receptor usage leading to distinct host and/or tissue tropisms and determining their antigenic profiles^12,13^.

A major challenge for the utilization of AAV vectors in the clinic is the presence of pre-existing neutralizing antibodies (NAbs) in a large percentage of the human population^14-16^. These antibodies originate from prior exposure to naturally circulating AAVs, primarily targeting the capsid, leading to vector inactivation and loss of treatment efficacy. As a result, patients having these antibodies, or exceeding defined antibody titer thresholds, are excluded from receiving these therapeutics^17,18^. One strategy to circumvent these antibodies is the engineering of capsid variants that are not bound by pre-existing NAbs. This approach requires the identification of the antibody epitopes on the capsid surfaces. For this purpose, the fragment antigen binding (Fab) domains of the immunoglobulins (Ig) are added to the capsid and the complex structure determined by cryo-EM. An Fab is composed of the light chain and the N-terminal half of the heavy chain. Each chain possesses three complementarity-determining regions (CDRs), which are responsible for recognizing specific antigens.

Initially, the resolutions of AAV capsid-antibody cryo-EM maps were low (∼5–20 Å)^8,19,20^. However, recent advancements of the cryo-EM technology have enabled the analysis of the interaction between the antibody and the capsid surface at atomic resolution^21^. The identified interacting residues form the basis for AAV capsid engineering efforts by rational design or by directed evolution^22^. Modifications of capsid aa have been shown to result in capsid variants capable of escaping the characterized antibodies *in vitro* and *in vivo*^19-21,23^. To date, this structural mapping approach has relied primarily on mouse monoclonal antibodies (mAbs) to simulate the human antibody response against the AAV capsids. A recent study obtained 21 individual human mAbs from three patients following Zolgensma^®^ treatment, which utilizes the capsids of AAV serotype 9^24^. The identification of their binding sites to the AAV9 capsid at low resolution showed that the majority of human mAbs bound to the 2-fold region of the capsid, whereas mouse mAbs preferentially bound to the 3-fold protrusions^19,24^.

The current study identified the interacting residues of these human mAbs using high-resolution cryo-EM. However, due to mAbs bound to the icosahedral symmetry axes, 3D-reconstructions of the complexes with imposed icosahedral symmetry resulted in “blurred” densities for the Fabs, preventing effective model building. To overcome the symmetry-mismatches, localized reconstructions with symmetry relaxation^25^ were utilized, enabling the deconvolution of the alternative binding modes of the Fabs. Utilizing this structural information, key residues in the AAV9 capsid were modified to generate a capsid variant escaping 18 of the 21 mAbs at high antibody titers.

This AAV capsid variant has the potential to expand the patient cohort treatable with AAV vectors to include patients that were previously excluded due to their pre-existing antibodies and opens the possibility of re-administration of the vector if such a need arises.

## Results

### Icosahedral averaging only resolves Fabs binding away from the symmetry axes

For a detailed understanding of the interaction between the human mAbs and the AAV9 capsids, high-resolution cryo-EM data was collected for each of the capsid-antibody complexes. Utilizing standard 3D-reconstruction protocols by imposing icosahedral symmetry using ∼43,000-366,000 particles resulted in cryo-EM maps at resolutions ranging from 1.88–3.27 Å (Tab.S1). For each map, the density contributed by the bound Fab was detected at the previously determined capsid regions^24^. At the 2/5-fold wall of the capsid Fab1-2 was resolved at a resolution of 2.61 Å (Fig.1A). In the map, the capsid, the variable heavy (V_H_) and light (V_L_) chains were well ordered and allowed the building of reliable models. In contrast, the constant regions of the Fabs were less ordered and not observed at a sigma (σ) threshold level of 2.0, likely due to a higher level of flexibility as they do not participate in the complex formation. Around the 5-fold axis, Fab1-6 and Fab2-7 were resolved at a resolution of 2.33 and 2.18 Å (Tab.S1) but adopting differential binding modes. While Fab1-6 bound closer to the 5-fold channel, Fab2-7 bound mostly to the depressed region surrounding the 5-fold channel, leaning towards the 2/5-fold wall (Fig.1B). In both maps the V_H_ and V_L_ chains were well ordered, enabling the building of atomic models, with the exception of some regions of the V_H_ chain of Fab1-6 located close to the 5-fold channel. The latter region was less structurally ordered and the built model of the Fab resulted in clashes with the neighboring, symmetry related Fabs. However, the disordered density observed for Fab1-6 was only minor in comparison to all 2-fold and 3-fold binding Fabs. The complexes with the two 3-fold binding Fabs, Fab1-1 and Fab3-4, were reconstructed to 2.62 and 3.27 Å resolution. In both cryo-EM maps, the capsid was well ordered. However, the density from the Fabs observed directly above the 3-fold symmetry axis was disorganized and could not be used to build atomic models (Fig.1C). This lack of structural ordering is the result of only a single Fab occupying the space at the 3-fold axis in one of three binding modes. Consequently, the bound Fab does not conform to the symmetry of the capsid and icosahedral averaging imposed during the 3D reconstruction blurred the density of the Fab. A similar situation was observed with the 2-fold binding Fabs. The complexes of the 16 Fabs binding to this region of the capsid were reconstructed to a resolution of 1.88–2.84 Å resolution (Tab.S1). In each map, the capsid was well ordered, while the densities for the Fabs above the 2-fold axis could not be interpreted for model building (Fig.1D). As for the 3-fold antibodies, only a single Fab bound to the 2-fold region, in two potential binding modes, resulting in distorted densities during icosahedral averaging.

**Figure 1.**
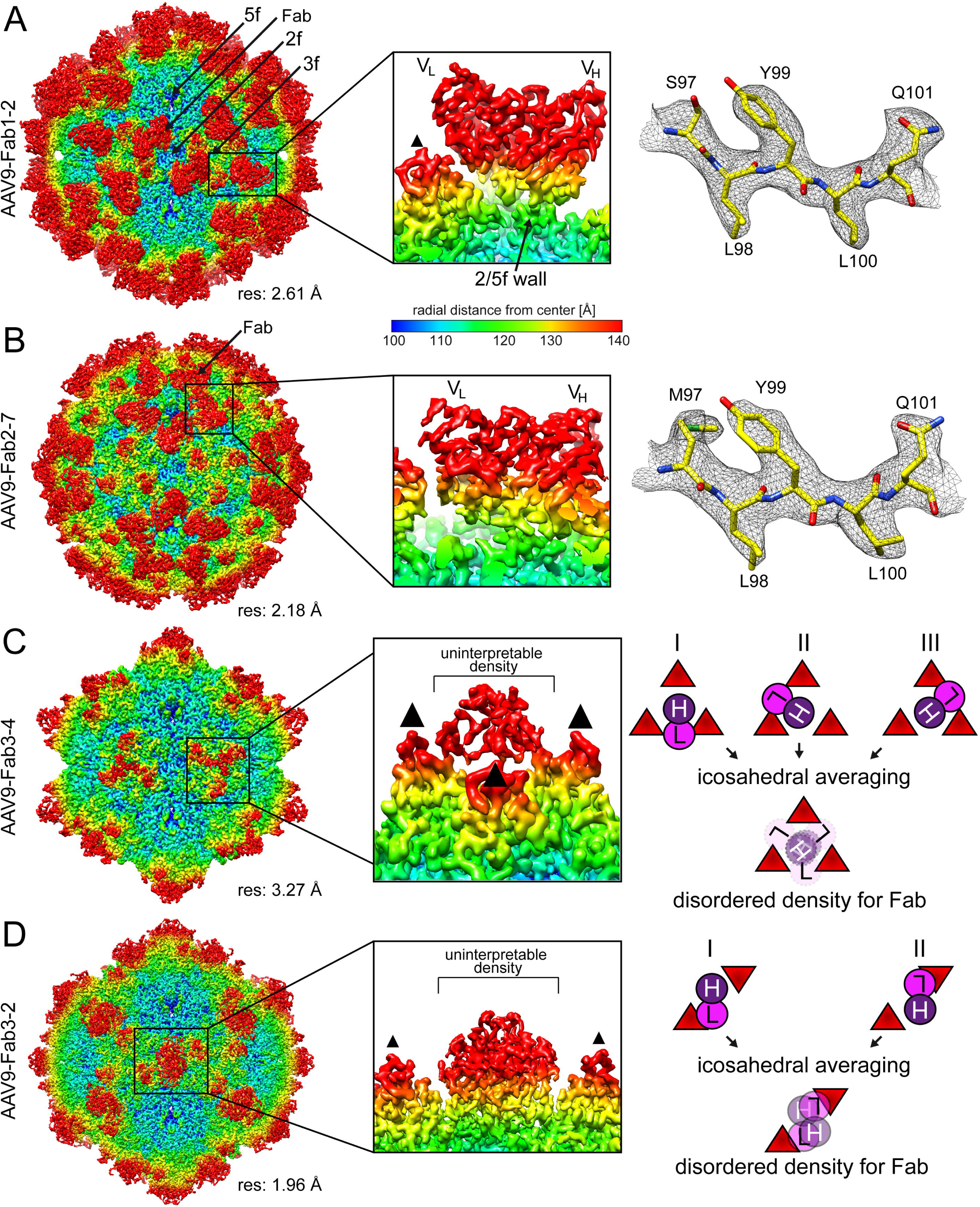
Icosahedral reconstruction of the AAV9-Fab complexes. A) Surface density map of the cryo-reconstructed AAV9-Fab1-2 complex. The view is down the icosahedral 2-fold axis, the map is contoured at a sigma (σ) threshold level of 2.0. The map is colored according to radial distance from the particle center (blue to red), as indicated by the scale bar. The icosahedral 2-, 3-, and 5-fold axes are indicated. The estimated resolution, determined at an FSC threshold value of 0.143, is shown below the map. The binding of the Fab on top of the 2/5-fold wall is shown as a close-up image in the center panel with the variable heavy (V_H_) and variab_l_e light (V_L_) chains labeled. To the right a representative stretch of amino acid residues modeled for the heavy chain of the Fab are shown inside the cryo-EM density map. The amino acid residues are labeled and shown as stick representations and colored according to atom type: C = yellow, O = red, N = blue, S = green. B) Depiction of the AAV9-Fab2-7 complex as in (A) with the Fab binding around the 5-fold symmetry axis. C) Surface density map of the AAV9-Fab3-4 complex as in (A). The Fab binds to the center of the 3-fold symmetry axis. Due to the imposed symmetry during the reconstruction process the density of the Fab is blurred and cannot be used to generate an atomic model. The illustration on the right gives a rationale for the blurring of a non-icosahedral Fab bound to an icosahedral capsid during icosahedral averaging. D) Depiction of the AAV9-Fab3-2 complex as in (C) with the Fab binding to the center of the 2-fold symmetry axis. All density map images were generated using UCSF-Chimera ^55^.

### Localized reconstruction resolves Fabs bound close to the symmetry axes

The visualization of asymmetric features in otherwise icosahedral structures, such as the Fabs binding near or at the capsids’ icosahedral symmetry axes, using traditional cryo-EM reconstruction procedures has been challenging as their information is lost due to averaging of the particle. However, standard asymmetric reconstruction is unfeasible here as the Fab binding modes are independent at each axis resulting in a vast number of structural classes (∼3^20^ possible permutations for the 3-fold binder)^26^. Thus, the localized reconstruction method was used in combination with symmetry relaxation to structurally characterize the 2-, 3-, or 5-fold region to reconstruct the Fabs not conforming to the icosahedral symmetry of the capsid^25-27^. For the sixteen 2-fold binding Fabs this strategy resulted in sub-particle maps reconstructed to 2.54 to 3.56 Å resolution (Tab.S1). The localized reconstructions with C2 symmetry relaxation confirmed that only a single Fab is bound at the 2-fold region (Fig.2A). In all maps, the V_H_ and V_L_ chains are well ordered enabling the building of atomic models. An exception was Fab2-6 with a well-ordered V_H_ chain but poorly ordered V_L_ chain, indicating that the V_L_ may not participate in capsid binding. Some maps also resolved the constant heavy (C_H_) and light chain (C_L_) of the Fabs such as Fab2-1 (Fig.2A). For the two 3-fold binding Fabs, Fab1-1 and Fab3-4, localized reconstruction also resolved the single Fab at the 3-fold axis at a resolution of 3.31 and 3.73 Å (Tab.S1 and Fig.2B). In the case of Fab1-1, the V_H_ and V_L_ chains are well ordered, whereas for Fab3-4 only the V_H_ chain is well-ordered, indicating that capsid binding is mainly mediated by V_H_. Lastly, for the 5-fold Fabs, the AAV9-Fab1-6 complex was processed with localized reconstruction, resulting in a 3.47 Å resolution map. In contrast to single Fabs binding at the 2- and 3-fold, the cryo-EM map revealed two bound Fabs with well-ordered V_H_ and V_L_ chains, surrounding the 5-fold axis in an opposing manner thereby avoiding clashes with Fabs bound at the nearest 5-fold related capsid monomer (Fig.2C).

**Figure 2.**
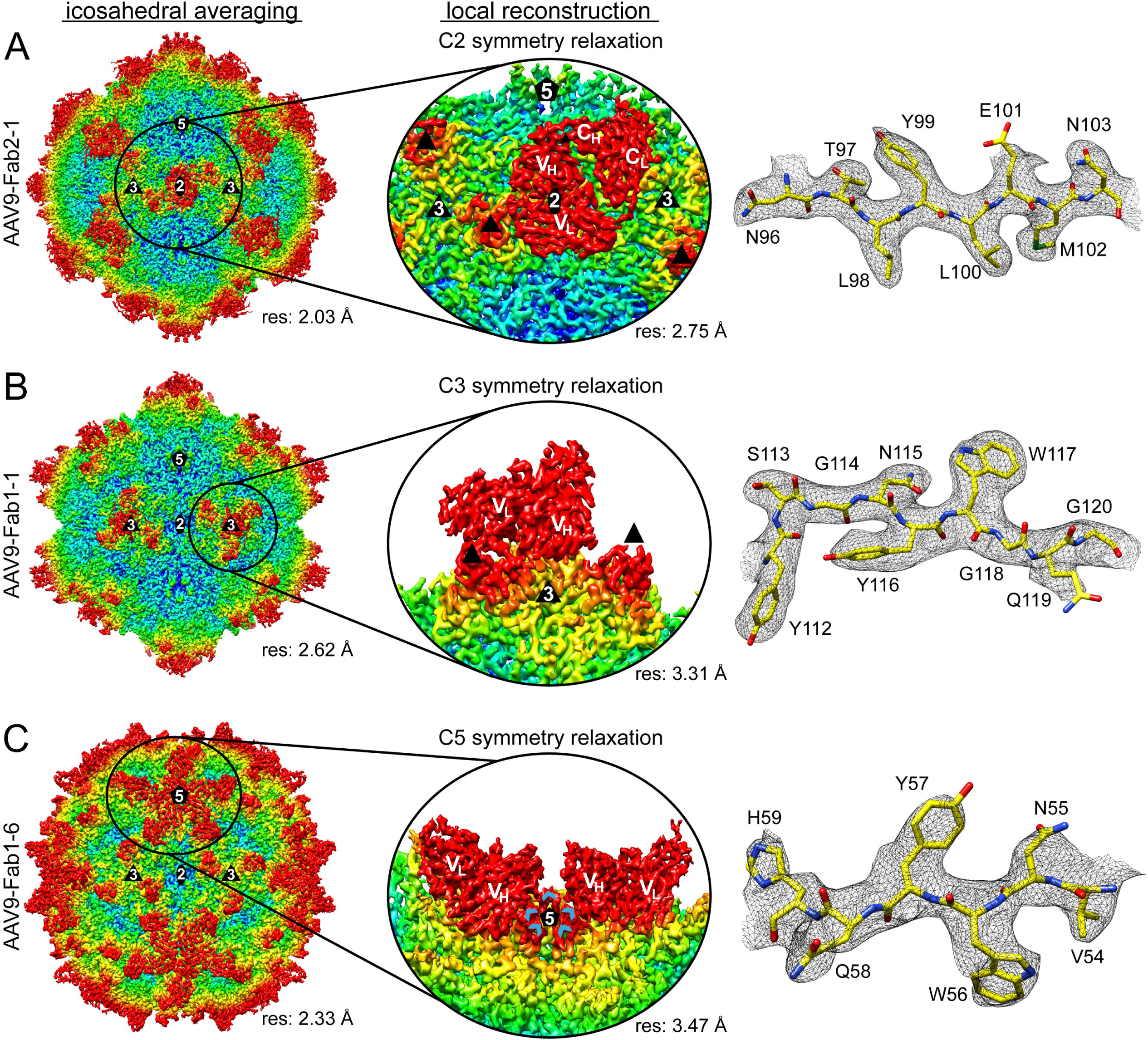
Localized reconstruction of the AAV9-Fab complexes. A) Surface density map of the icosahedral-reconstructed AAV9-Fab2-1 complex colored as in Fig. 1. The icosahedral 2-, 3-, and 5-fold axes are indicated, and the estimated resolution is shown below the map. To the right a density map of the 2-fold region is shown derived from localized reconstruction with C2 symmetry relaxation. The variable/constant heavy (V_H_/C_H_) and vari_a_bl_e_/constant light (V_L_/C_L_) chains labeled. The 3-fold protrusions are indicated as black triangles. A representative stretch of amino acid residues modeled for the heavy chain of the Fab are shown inside the localized reconstructed map. The amino acid residues are labeled and shown as stick representations and colored according to atom type: C = yellow, O = red, N = blue, S = green. B) Depiction of the AAV9-Fab1-1 and C) Fab1-6 complex as in (A). For the localized reconstruction of the 3-fold region C3 symmetry relaxation and the 5-fold region C5 symmetry relaxation were applied, respectively.

### The 2-fold binding Fabs exhibit four distinct binding modes

The atomic models for the Fabs allowed the comparison of their binding patterns to the AAV9 capsid. The sixteen Fabs binding to the 2-fold axis could be structurally sub-classified into 4 groups (Fig.3A). In group A, containing Fab2-1, 2-3, 2-4, 2-6, 3-1, 3-2, and 3-5, the V_H_ chain of the Fab binds perpendicular to the 2-fold axis with its CDRs entering the depression. Compared to V_H_, the V_L_ chain is shifted toward the 2/5-fold wall and rotated approximately 90° with its CDRs binding to the side of the 3-fold protrusions. The group B Fabs, Fab1-5, 2-2, 3-6, and 3-7, are rotated ∼20° relative to group A, shifting the V_L_ chain away from the side of the 3-fold protrusions. In group C, comprising of Fab1-3, 1-4 and 2-5, it is the V_L_ chain that is situated above the 2-fold axis. The V_H_ chain is shifted towards the 2/5-fold wall binding to the side of the 3-fold protrusions with CDR1/2, whereas the long CDR3 loop enters the 2-fold depression. This binding mode resembles the group A Fabs with the position of the heavy and light chains swapped. However, in both groups the CDR3 loop of the V_H_ chain occupies approximately the same space in the 2-fold depression. In group D, Fab1-7 and 3-3, the V_H_ chain is located on top of the 2-fold axis with CDR1 and CDR2 entering the 2-fold depression while CDR3 binds to the side of the 3-fold protrusions. Unlike for group A, the V_L_ chains are elevated higher from the capsid and their CDRs making contact to the top of the 3-fold protrusions. To investigate whether the grouping can be correlated to the Fabs aa sequence, phylogenetic trees for the V_H_ and V_L_ chains were generated. In the analysis the V_H_ chains did not form clusters for the individual Fab groups with the exception of group D (Fig.S1A). A common feature of the group D Fabs is their much shorter CDR3_H_ loop (Fig.S1B), possibly because it does not enter the 2-fold depression. In contrast, the group A Fabs possess on average the largest CDR3 loop for the V_H_ chain. This is probably the result of the chain slightly shifting towards the 2/5-fold wall and the need to still enter the 2-fold depression. The remaining five CDRs of the Fab do not show significant length differences as previously reported^24^. The phylogenetic analysis of the V_L_ chains showed a clustering of group B and most of group A Fabs (Fig.S1C) suggesting a potential role of the light chain for the binding mode of the Fab.

**Figure 3.**
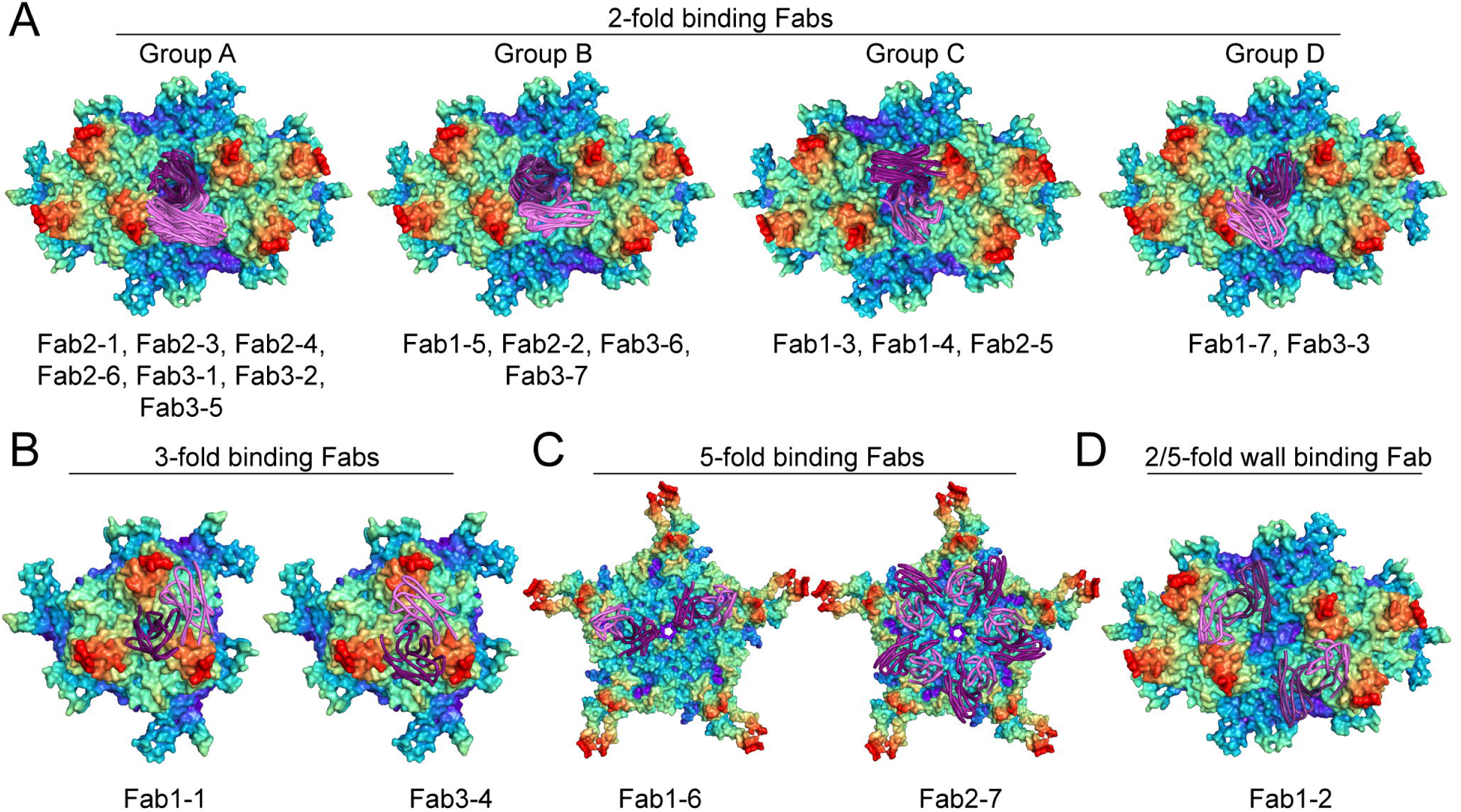
Binding mode of the Fabs to the AAV9 capsid. A) Radial-colored surface representations of the AAV9 capsid are shown for double-trimers with the 2-fold symmetry axis in the center. The structures of the V_H_ (purple) and V_L_ (pink) chain are shown as ribbon diagrams. The 2-fold binding Fabs are classified into 4 groups (A, B, C and D) based on their interaction to the AAV9 capsid. The Fabs belonging to the groups are listed below each surface representation. B) Surface representations of AAV9 capsid trimers showing the binding of Fab1-1 and Fab3-4. C) AAV9 capsid pentamer showing the binding of Fab1-6 and Fab2-7. D) Surface representation as in A for Fab1-2.

The Fabs, 1-1 and 3-4, are located differently in the 3-fold region. While the V_H_ chain of Fab1-1 is positioned directly above the 3-fold symmetry axis, V_H_ of Fab3-4 is located further away from the 3-fold axis between two protrusions (Fig.3B). Despite their different locations, the CDR3 loop of both Fabs enter the depression at the center of the 3-fold axis. To achieve this, the CDR3 of Fab3-4 is three aa longer. The V_L_ chains of both Fabs are situated between the protrusions, with the V_L_ chain of Fab1-1 being shifted further away from the 3-fold axis.

The two 5-fold Fabs differ in their binding behavior (Fig.3C). While the V_H_ chain of Fab1-6 is located close to the 5-fold channel, V_H_ of Fab2-7 is located closer to the 2/5-fold wall. However, with its ten aa longer CDR3_H_ loop, Fab2-7 is capable of interacting with the 5-fold channel. Furthermore, the V_L_ chains of both Fabs interact with the AAV9 capsid in approximately the same region. Lastly, Fab1-2 binds to the 2/5-fold wall with its V_H_ chain while the V_L_ chain is situated between the 3-fold protrusions (Fig.3D).

### The cryo-EM maps reveal the capsid-antibody contacts

The high resolution of the complex maps allowed the building of models for the Fabs and the capsid (Fig.4A). Of particular interest were the CDRs of the Fabs that mediate the interaction to the capsid. The CDRs of the heavy chains (Fig.4B) and light chains (Fig.4C) exhibited well-resolved densities for the aa side chains enabling the identification of the capsid-antibody contacts. Hydrogen bonds, Van-der-Waals contacts and hydrophobic interactions up to a distance of 3.5 Å, in addition to salt bridges up to 4 Å were considered as contacts (Fig.4D). In some cases, the interactions involved intermediate densities, such as in between N78 (Fab2-1) and T492, that were interpreted as water molecules. Other interactions involved bivalent cations, that were interpreted as calcium ions, e.g. between E125 (Fab2-3) and a series of AAV9 residues (Fig.S2). All contacts for the Fabs from the three patients are listed in table S2, S3, and S4. Overall, Fab1-3 had the lowest number of contacts with the AAV9 capsid, followed by Fab1-1. The Fabs with the highest number of contacts all belong to group 2 of the 2-fold binders. Moreover, all Fabs interact with multiple VPs on the AAV9 capsid surface, which confirms that all mAbs exclusively detect intact capsids (Fig.S3).

**Figure 4.**
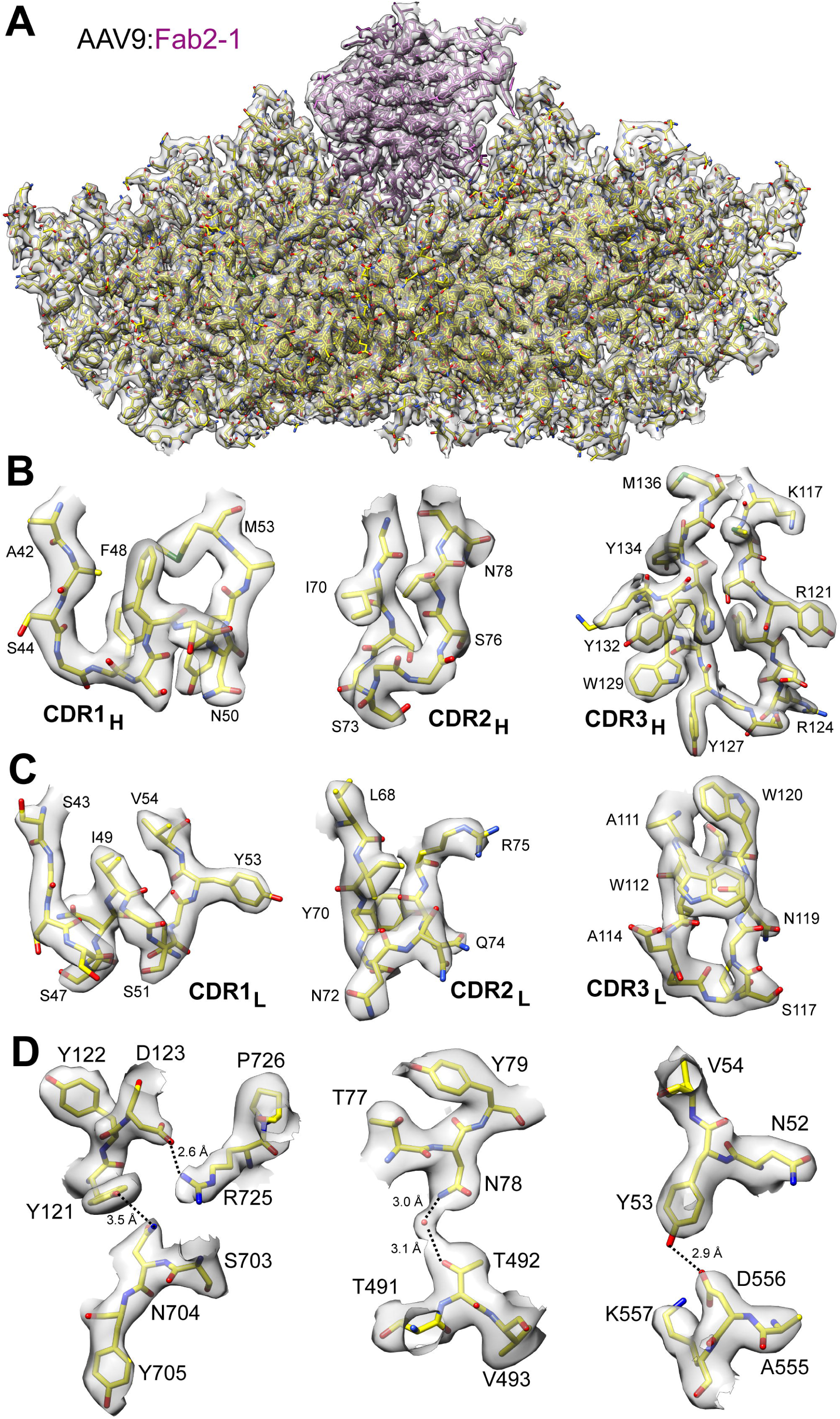
Determining the contacts of Fab2-1 to the AAV9 capsid. A) Modeled CDR1-3 of the heavy chain inside the cryo-EM map. The contacts to the AAV9 capsids including the distance of the interactions are listed below B) Shown inside the cryo-EM map is the fit of the AAV9 capsid (yellow) and Fab2-1 (purple) models. C) The modeled CDR1-3 of the light chain inside the cryo-EM map is shown as in (A). D) Exemplary contacts of the Fab to the AAV9 capsid are shown.

For the subsequent capsid engineering, the VRs and the frequency of the contacts on the capsid surface were analyzed. The epitopes for the sixteen 2-fold Fabs comprise VR-III to - VII, and -IX of the AAV9 capsid (Fig.5A-C). However, only group B Fabs contact VR-III. On the aa level, all 2-fold-Fabs contact D532 and Y706. Other frequently contacted residues are T491, T492, R533, D556, N562, N704, and Y705, which are clustered around the 2-fold symmetry axis (Fig.5D). The contact residues for Fab1-1 and Fab3-4 cluster around the 3-fold symmetry axis and involve aa in VR-V and -VIII, including T584 and Q588 (Fig.S4). For the 5-fold binding Fabs, the contacted aa are more spread out and are located in VR-I, -II, -VII, the HI-loop, and VR-IX. Finally, the contacts of Fab1-2 are distributed to VR-I, -III, -VI, and VIII.

**Figure 5.**
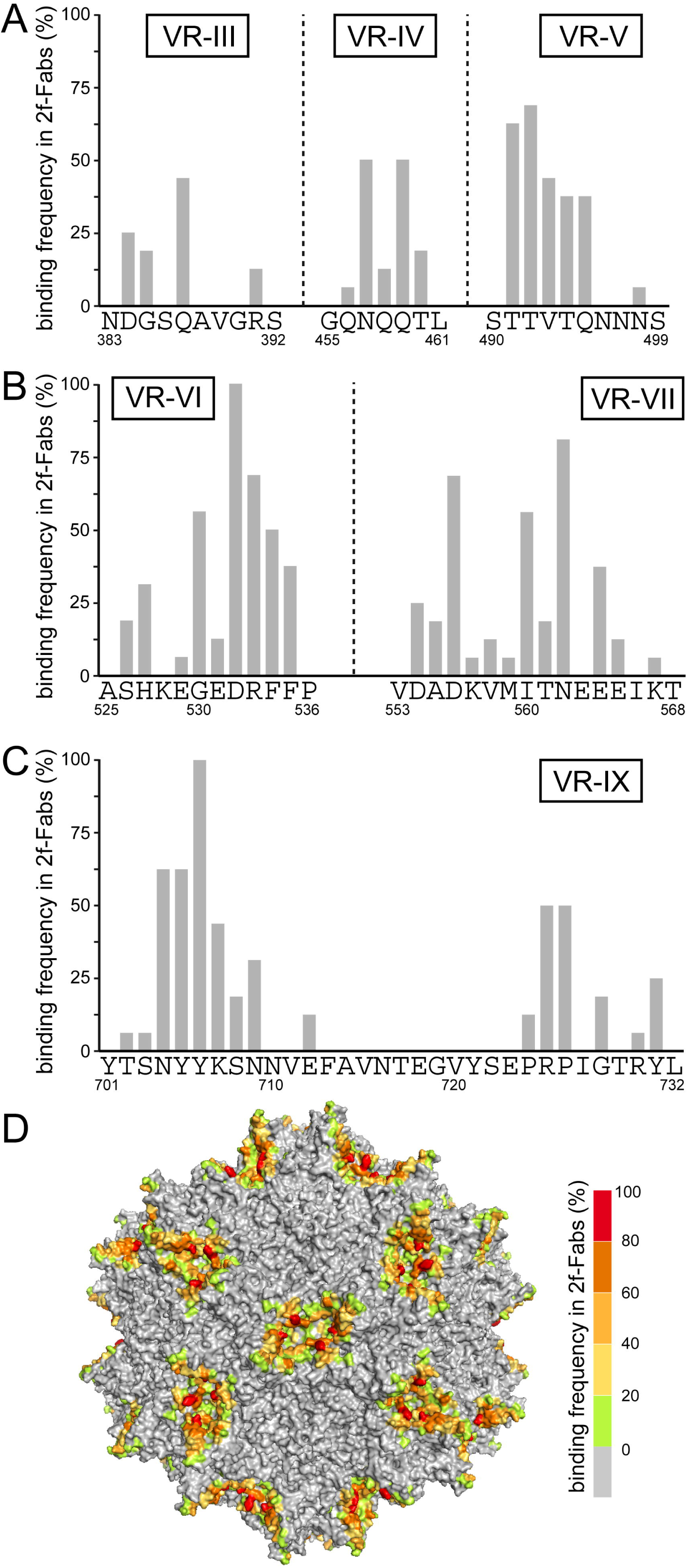
Contact frequency of the 2-fold binding Fabs to the AAV9 capsid. **A)** The percentages of contacted capsid residues within VR-III, -IV, -V, **B)** -VI, -VII, and **C)** VR-IX are shown for all 2-fold binding Fabs. **D)** An AAV9 capsid surface representation with the binding frequency for any 2-fold binding Fabs is displayed which is colored according to the bar on the right.

### The Fabs induce conformational changes of the AAV9 capsid

The cryo-EM maps allowed the detection of induced conformational changes of the AAV9 capsid upon antibody binding. The binding of most Fabs induced structural changes at the Fab binding site. For Fab1-2, binding to the 2/5-fold wall, the antibody caused a rearrangement of VR-I, shifting the position of some aa by up to 11 Å (Fig.6A). Fab1-6 pushes the loop forming the 5-fold channel (VR-II) ∼5 Å inwards restricting the diameter of the channel (Fig.6B). The adjacent DE-loops not bound by the Fab adopt the canonical loop conformation of the AAV9 capsid in absence of the Fab. A similar observation was made for Fab1-1 which uses Q588 as a contact residue by altering its side chain orientation in one of the 3-fold protrusions (Fig.6C) whereas the other two 3-fold protrusions adopt the unbound conformation. Fab1-7 and 3-3, belonging to group D of the 2-fold binders interact with the apex of the 3-fold protrusions, resulting in shifts of VR-IV of up to 8 Å (Fig.6D). Furthermore, for all the 2-fold Fab-complexes movements occurred in VR-IX near the 2-fold axis. While, for AAV9 capsids in absence of Fabs the residues 704-707 adopt the same conformation in both symmetry-related VR-IX loops surrounding the 2-fold axis, there are numerous variations of side-chain orientations in the different Fab complex structures (Fig.S5). These changes are likely caused by the Fabs contacting these residues or by the Fab moving the capsid side chains into a more favorable position for antibody binding. Interestingly, none of the complex structures adopt the same conformation for residue range 704-707 in both symmetry-related VR-IX loops.

**Figure 6.**
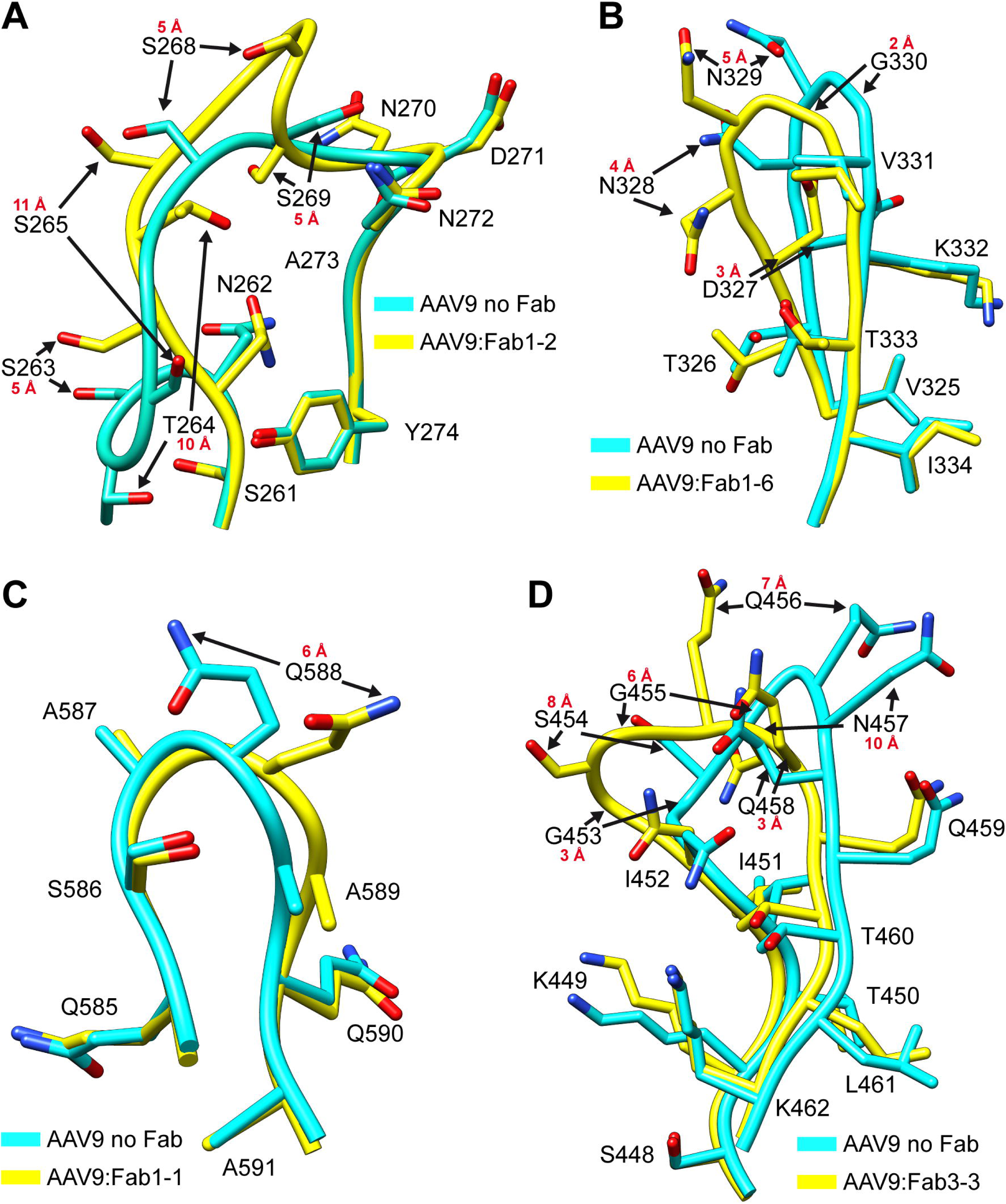
Fab induced structural rearrangement of the AAV9 capsid. A) Binding of Fab1-2 to the AAV9 capsid leads to an alternative conformation of VR-I. The conformation in absence of the antibody (cyan) and in presence of Fab1-2 (yellow) are shown as ribbon diagrams with amino acid side chains. Movements of ≥ 3 Å between the indicated amino acids are indicated. B) Depiction as in (A) for VR-II upon binding of Fab1-6, C) VR-IV upon binding of Fab3-3, and D) VR-VIII upon binding of Fab1-1.

### Amino acid changes at the Fab contact sites enable antibody escape

The structural data of the AAV9-Fab complexes guided the development of AAV9 capsids with antibody escape phenotypes. Amino acid substitutions of the identified contact residues were selected either by disrupting critical interactions of the Fab with the capsid or by introducing steric clashes, preventing antibody binding (Fig.S6). The newly generated capsids were analyzed for their productivity and infectivity to exclude non-viable variants. Subsequently, these variants were tested for their ability to escape the mAbs. Successful aa substitutions were combined to generate an AAV9 capsid variant capable of evading multiple antibodies. The newly engineered variant of this study is composed of six aa changes, T491R, D556P, N562Y, T582Q, Q588R, and Y706D, henceforth called hAEV6 (human antibody escape variant 6) (Fig.7A). In native immuno-dot blots this variant was not recognized by any of the 2-fold and 3-fold antibodies, amounting to 18 out of the 21 human mAbs (Fig.7B). As none of the aa substitutions target the 5-fold and 2/5-fold wall binding mAbs, hAEV6 was still detected by Fab1-2, Fab1-6, and Fab2-7. Several variants for these mAbs were developed but either failed to escape the antibodies or resulted in defective capsids (Tab.S5). The hAEV6 capsid does not change previously identified residues of the described galactose binding pocket but one aa (Q588R) of the AAVR binding interface^28,29^ and surprisingly showed a ∼7.5-fold higher transduction efficiency vs. AAV9 (Fig.7C). When challenged with the pooled 18 mAbs binding to the 2- and 3-fold region hAEV6 transduction is not inhibited, even in the presence of high mAbs concentrations (Fig.7D). In contrast, AAV9 lost its ability to transduce at 100-1000-fold lower mAbs titers. Utilizing pooled mAbs including all 21 mAbs did not alter the neutralization curve for AAV9 but neutralized hAEV6 at the two highest antibody concentrations. However, compared to AAV9, hAEV6 neutralization is shifted to levels where ∼10-fold higher mAbs concentrations are required for equivalent neutralization. This mimics the endpoint titer determined from human sera of six Zolgensma patients which were ∼13-fold lower for the hAEV6 compared to AAV9 capsid (Fig.7E).

**Figure 7.**
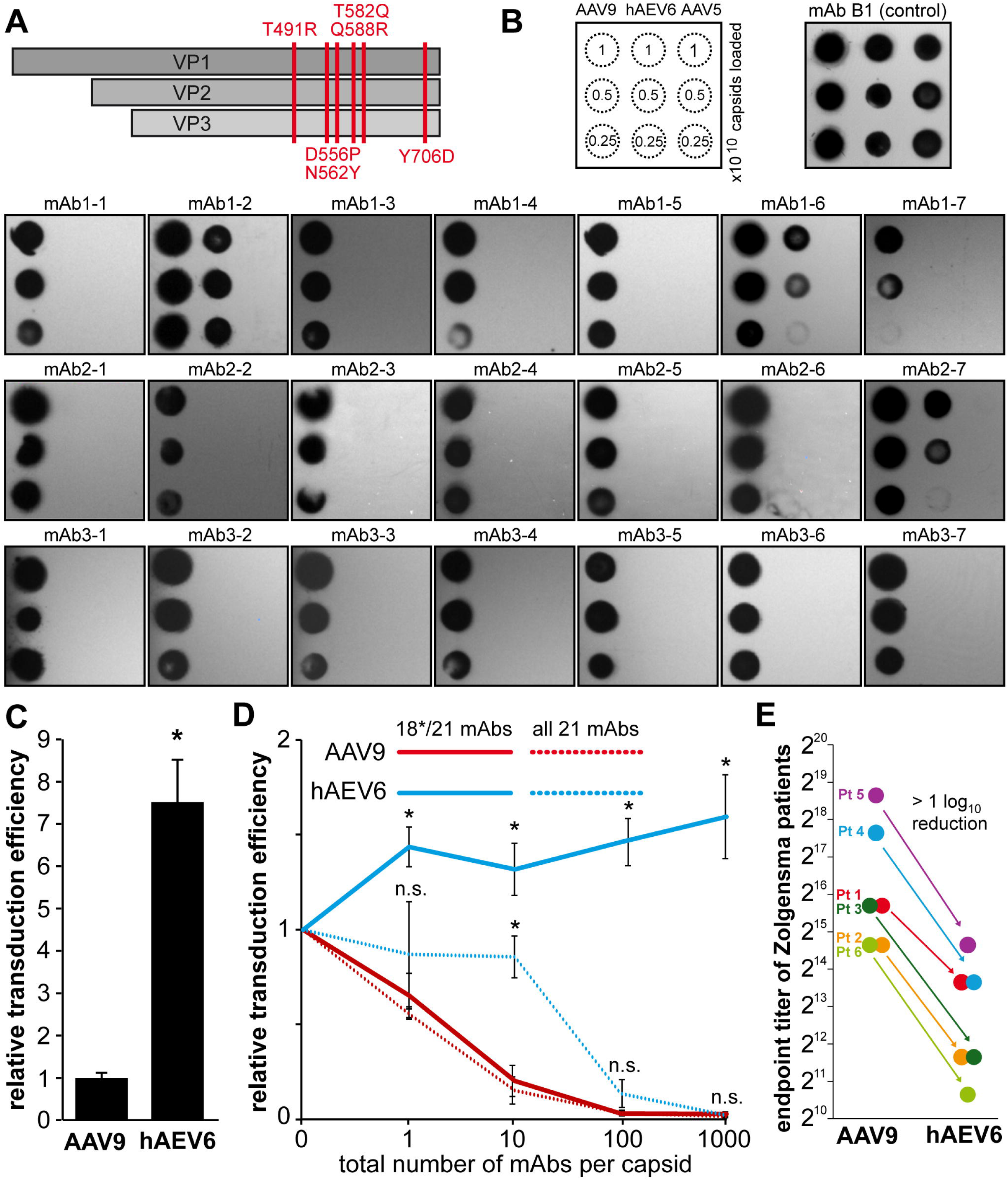
Generation of an antibody escape variant. **A)** Schematic showing the amino acid substitutions of hAEV6 **B)** Native dot blots are shown using mAb B1 as a loading control and the 21 human mAbs against the capsids of AAV9, hAEV6, and AAV5. The hAEV capsid variant escapes 18 of the 21 human antibodies, with the exception of mAb1-2, 1-6, and 2-7. **C)** Transduction analysis showing that hAEV6 displays enhanced transduction efficiency relative to AAV9. **D)** Transduction of hAEV6 is maintained in the presence of a pooled mAb solution which contained equal amounts of the 18 mAbs that the aa substitutions of hAEV6 are targeted against or equal amounts of all 21 mAbs. All experiments have been performed in triplicates (n=3). Asterisk indicates p< 0.01. **E)** Sera from Zolgensma-treated infants (n = 6) were analyzed to determine the anti-AAV9 or anti-hAEV6 IgG endpoint titers. The titers of individual patients (Pt) are colored and the average reduction of the endpoint titer from AAV9 to hAEV6 calculated.

## Discussion

To this date, all approved AAV biologics utilize the capsids of naturally occurring AAVs^30,31^ circulating in the human population, resulting in worldwide seroprevalence rates of 20-80% against different AAV serotypes^14,32^. While antibodies generated by the adaptive immune system are generally favorable to eliminate a pathogen from the host and to prevent future re-infections of the same agent, they are detrimental to the success of AAV-mediated gene therapy. To circumvent these antibodies, future AAV biologics may need to utilize capsids where most if not all antigenic epitopes of the natural-circulating AAVs have been eliminated. These engineered capsids will not prevent future immune responses upon vector administration but will expand the patient cohort treatable with AAV vectors that previously were precluded due to their pre-existing antibodies. In the past 12 years, the antibody responses against the AAV capsids were primarily simulated with mouse mAbs^13,20,33,34^. However, our recent study characterizing 21 anti-AAV9 capsid mAbs from human patients indicated that human and mouse antibodies may exhibit differential binding characteristics to the AAV capsids^24^. Prior studies showed that 66% of mouse mAbs bound to the 3-fold region whereas 75% of human mAbs bound to the 2-fold region of the AAV9 capsid^19,33^. This observation was unusual because none of the 22 mouse mAb developed against AAV capsids to date have been observed to bind directly to the 2-fold region of any AAV capsid^13^. However, it should be noted that some antibodies, such as 4E4, ADK4, and ADK9 bridge across the 2-fold axis without binding to the depressed region of the capsid^12,13,19^. Consequently, the 2-fold region was considered to be the least antigenic region of the AAVs, potentially due to the Fab being sterically unable to enter the 2-fold depression. A possible explanation that allows human antibodies to contact the residues inside the depression are their longer CDRs compared to mouse antibodies, which were reported to be 15.5±3.2 and 11.5±1.9 aa for the human and mouse CDR3 ^35^. The lengths of the CDR3_H_ loops of the 2-fold binding mAbs in the present study were even longer, ranging from 14-24 aa. For comparison the CDR3_H_ loop of mouse mAbs with known sequences binding to AAV capsids have a length of 12 (A20) and 11 aa (HL2476)^21,34^. However, the length alone does not explain the preference for the 2-fold region. A comparison of previously determined antibody-antigen interactions indicated that there is a tendency for hydrophobic aa to reside in the center of an antibody epitope, flanked by charged aa^36^. The same study also analyzed the aa preference within epitopes, with the top 4 overrepresented residues being tryptophan, tyrosine, methionine, and histidine, whereas valine, alanine, leucine, and cysteine were the most underrepresented residues. These conditions are met at the 2-fold region of the AAV9 capsid and may be the driver for the antigenic hotspot (Fig.S7).

The HL2476-Fab bound to the capsid of AAV5 is currently the only capsid-antibody complex determined at high-resolution (3.1 Å)^21^. This Fab was found to bind to the 3-fold protrusion, a total of 60 times. As such, the Fab followed the icosahedral symmetry of the capsid, enabling the modeling of the complex. The remaining AAV-Fab complexes have been determined at resolutions ranging from 4.1-23 Å^13,19,20,37^. If high-resolution structure determinations using cryo-EM and conventional icosahedral reconstruction would have been attempted for antibodies binding near the 2-fold (e.g. 4E4, ADK4, ADK9), at the 3-fold (e.g. 5H7, PAV9.1), or around the 5-fold (e.g. HL2372) the structural information for the Fabs would have been lost due to icosahedral averaging of the particle, similarly as observed in this study. Binding of Fabs directly at one of the symmetry axes precludes another Fab from binding in one of the other symmetry-related binding modes. As a result, Fabs binding at the 2-, 3-, and 5-fold axis are limited to 30, 20, and 12 binding events to the icosahedral capsid, respectively. The 5-fold binding Fabs in this study bind offset from the symmetry axis. However, due to the fact that Fab1-6 would be clashing with Fabs bound to its symmetry-related neighboring binding sites, this Fab is limited to 24 binding events to the icosahedral capsid. In contrast, no clashes were observed for Fab2-7, enabling the binding of up to 60 Fabs to the capsid. In order to resolve symmetry-mismatches, localized reconstruction methods have been developed to structurally characterize patches of the capsid for ligands not conforming to the icosahedral symmetry^25-27^. Prior to this study, asymmetric features of AAV capsids have not been analyzed, except for a study of AAV2 in complex with AAVR determined by cryo-electron tomography at 20-30 Å resolution and a recent study that used a similar strategy to localized reconstruction for an AAV9 capsid variant binding to its receptor carbonic anhydrase IV^39^.

For some of the capsid-Fab complexes, ordered water molecules were observed in the binding interface. This has been previously observed for antigen-antibody structures and suggested to mediate and stabilize the antigen-antibody association^40^. Additionally, at least two Fabs showed the presence of a bivalent cation in the binding interface. Previous studies have shown that some antibodies depend on the presence of calcium ions for the binding of its antigen^41,42^. Future characterizations of the mAbs will confirm if capsids treated with EDTA will prevent antibody binding or reduce their binding affinity.

Conformational changes in the antibody following antigen binding, especially in the CDRs, have been widely reported^43-46^, however, studies observing structural changes in the antigen upon antibody binding, as observed here, are less common. However, a recent study showed that binding of antibodies to SARS-CoV-2 variants induced conformational changes in the viral spike protein^47^. These observations are only detectable at sufficient resolution. For the AAV5-HL2476 complex, only minor shifts up to 1.1 Å in VR-V were observed which was the main contact region of the Fab^21^. Similarly, conformational changes in the capsid of a distantly related parvovirus (canine parvovirus), upon binding with Fab14 were also minor with a maximum displacement of the main chain by 1.9 Å^48^. For the Fabs in this study conformational changes of the main chain varied from minor (Cα-RMSD ≤2 Å) for the 3-fold and most 2-fold binding Fab; medium changes (Cα-RMSD 2-5 Å) for the 5-fold Fabs; to significant remodeling of the surface loops (Cα-RMSD 5-11 Å) for the group D 2-fold Fabs and Fab1-2 binding to the 2/5-fold wall. Additionally, many aa side chains in the epitopes adopt alternative rotamer conformations. This was most pronounced for the 2-fold binding Fabs leading to various conformations of aa 704-NYYK-707. The 2-fold region of the AAV capsids has been suggested to play a role in viral transcription^49^ and may require similar dynamic aa configurations for this function.

Side chain orientation shifts can be caused by interactions to the Fab making these residues prime targets for capsid engineering, exemplified by the Q588R variant which escapes Fab1-1. Previous AAV9 variants developed for the escape of PAV9.1 swapped the residues aa586-590 with the corresponding residues from other AAV serotypes^33^ that may also escape Fab1-1. However, to prevent the reintroduction of antigenic epitopes from other AAVs, a more targeted approach was conducted, facilitated by the higher resolution structure, and Q588 substituted to a residue not found in other AAV serotypes. The AAV9 variants developed to escape the other mouse mAbs, S454A and P659K^19^, had no impact on the human mAbs but vice-versa Q588R also allowed escape from ADK9. The AAV9-hAEV6 capsid was able to prevent neutralization from 18 of the 21 human mAbs, comprising all AAV9-specific antibodies^24^. To date, no single aa substitution capsid variant was identified capable of escaping the mAbs binding to the 5-fold region and the 2/5-fold wall which cross-react to multiple AAV serotypes, including non-primate AAVs^50^. Likely, multiple aa substitutions will be required to disrupt the binding of these antibodies. While none of the changes in the hAEV6 variant targeted these antibodies, surprisingly, a shift in the neutralization assay was observed, allowing this capsid variant to tolerate ∼10-fold higher antibody concentrations before being neutralized. This effect could be associated with the higher transduction efficiency of the hAEV6 variant. Future studies will further characterize this variant to verify if the tissue tropism is comparable to AAV9 *in vivo*. If successful, this variant has the potential to be an alternative for AAV9 vectors. Currently, the exclusion criteria for receiving Zolgensma are anti-AAV9 antibody titers >1:50. Thus, a ∼10-fold higher tolerance against anti-AAV9 antibodies will lead to a significantly lower rejection rate of patients in need of this life-saving medication and possibly also allows redosing of patients previously treated with AAV9 vectors.

## Acknowledgements

The authors would like to thank the late Dr. Mavis Agbandje-McKenna for her pioneering studies of AAV capsid structures. The authors also thank the UF-ICBR electron microscopy core for access to electron microscopes utilized for cryo-electron micrograph screening (RRID:SCR_019146). High-resolution cryo-EM data collection was performed at the Stanford-SLAC Cryo-EM Center (S^2^C^2^), which is supported by the National Institutes of Health Common Fund Transformative High-Resolution Cryo-Electron Microscopy program (U24 GM129541). The content is solely the responsibility of the authors and does not necessarily represent the official views of the National Institutes of Health.

## Funding

The study was funded by an NIH grant R01 NIH GM082946 (to RM), a National Health and Medical Research Council of Australia grant (APP2004320 to IEA and GJL; and APP1194940 to MAF)), the Rebecca Cooper Foundation (PG2019449 to GJL) and a Research Council of Finland grant (348021 to JTH).

## Online Methods

### Cell culture

HEK293 cells were maintained adherent in Dulbecco’s Modified Eagle Medium (DMEM) (Thermo Fisher, Waltham, MA), supplemented with 10% heat-inactivated fetal calf serum and 100 units of penicillin/ml and 100 μg of streptomycin (Caisson Laboratories, Smithfield, UT) at 37°C in 5% CO_2_.

### Site-directed mutagenesis

The R2V9 plasmids containing AAV2 *rep* and AAV9 *cap* gene served as the template for site-directed mutagenesis PCR reactions. For each mutant, complementary PCR primers were designed that contained the desired mutation, which was flanked on both sides by 10 to 15 homologous base pairs. Primers were ordered from Sigma-Aldrich (Houston, TX) and used in PCR amplification reactions using a C1000 Touch™ thermal cycler (Bio-Rad, Hercules, CA) and *Pfu* Ultra high-fidelity DNA polymerase (Agilent, Santa Clara, CA). PCR products were incubated at 37℃ for 1 h with *Dpn*I restriction enzyme (NEB, Ipswich, MA) to degrade the methylated template plasmid. The reactions were then transformed into Top10 competent cells (Thermo Fisher), which were cultured on LB-ampicillin selective media and further amplified to isolate the plasmid. Clones were submitted for Sanger sequencing (Genewiz, South Plainfield, NJ) to verify the introduced mutations.

### Recombinant AAV production and purification

Recombinant AAV vectors with a packaged luciferase gene were produced by triple transfection of HEK293 cells at a confluency of ∼80-90%, utilizing pTR-UF3-Luciferase, pHelper (Stratagene), and R2V9 or variants thereof. The transfected cells were harvested 72 h post transfection, washed with phosphate-buffered saline (PBS; 137 mM NaCl, 2.7 mM KCl, 100 mM Na_2_HPO_4_, 2 mM KH_2_PO_4_), the cells pelleted and resuspended in PBS with 1 mM MgCl_2_ and 2.5 mM KCl. The resuspended cells were subjected to three freeze-thaw cycles (-80°C to 37°C) and subsequently incubated with 125 units/mL benzonase for 1 h at 37°C before centrifugation at 10,000 × g for 15 min to pellet the cell debris. AAV9 vectors or variants thereof were purified by Capture Select AAV9 affinity chromatography as previously described ^51^. The purity of the AAV capsid samples was analyzed by SDS-PAGE. For the determination of the titer of vector genome-containing capsids 5 μL aliquots were incubated with proteinase K at 56°C for 2 h, and the released vector genomes purified by a PCR purification kit (Invitrogen). Subsequently, a quantitative PCR was conducted in a Bio-Rad MyiQ2 Thermocycler instrument (Bio-Rad) using primers amplifying a 146 bp segment of the luciferase gene and the SYBR Green Master Mix (Bio-Rad).

### Native dot immunoblot analysis

Native AAV9 capsids were adsorbed onto nitrocellulose membranes (Bio-Rad, Hercules, CA) in a dot blot manifold (Schleicher and Schuell, Dassel, Germany). For loading controls and to confirm whether the capsids are exclusively detected in their native state, AAV9 capsids were heated at 100°C for 5 min prior to adsorption to the membrane. Excess fluid was drawn through the membrane by vacuum filtration. The membrane was removed from the manifold and blocked with 6% milk in PBS, pH 7.4 for 1 h. The human mAbs or mAb B1 were applied to the membrane at a concentration of ∼ 1 µg/ml in PBS with 6% milk, 0.1% Tween-20 and incubated for 1 hr. The membrane was then washed with PBS and an anti-human-HRP (Abcam) or anti-mouse-HRP secondary antibody (Cytiva) applied at a dilution of 1:50000 or 1:3000 in PBS/6% milk, respectively, and incubated for 1 h. The membrane was washed with PBS and then Immobilon™ Chemiluminescent Substrate (Millipore, Darmstadt, Germany) was applied to the membrane and the signal detected on X-ray film.

### AAV transduction and neutralization assay

Purified AAV9 vectors or AAV9 variants were used to infect HEK293 cells seeded on 24 well-plates at a MOI (multiplicity of infection) of 100,000. After 48 h, cells were lysed and luciferase activity assayed using a luciferase assay kit (Promega, Madison, WI) as described in the manufacturer’s protocol. Uninfected cells of the same plate were used as a negative control. For neutralization assays the vectors were pre-incubated for 30 min at 37°C with either purified mAbs, or PBS as the negative control at variable ratios relative to the capsids prior to infection. The transduction efficiency was calculated as a percentage to AAV vectors in the absence of antibodies.

### Cryo-EM sample preparation and data collection

Purified AAV9 vectors were mixed with Fabs at a ratio of ∼2 Fabs per potential VP binding site in the capsid, resulting in a final ratio of ∼1:120 (capsid to Fab). Three and a half microliters of the AAV9-Fab complex samples were applied to glow-discharged Quantifoil copper grids with 2 nm continuous carbon support over holes (Electron Microscopy Sciences), blotted, and vitrified in liquid ethane using a Vitrobot Mark 4 (FEI) at 95% humidity and 4°C. The grids were screened for particle distribution and ice quality with an FEI Tecnai G2 F20-TWIN microscope (FEI Co., Hillsboro, OR, USA) operated under low-dose conditions (200 kV, ∼20 e^−^/Å^2^). Images were collected on a Gatan UltraScan 4000 CCD camera (Gatan, Inc., Pleasanton, CA, USA). High-resolution data for the AAV9-Fab complexes were collected at the Stanford-SLAC Cryo-EM Center (S^2^C^2^), using a Titan Krios (FEI) electron microscope operated at 300 kV, equipped with a Falcon 4 direct electron detector (Thermo Fisher). A total of 50 movie frames were collected per micrograph at a total electron dose of ∼50 e^−^/Å^2^, a defocus range of -0.8 to -2.1 µm, a pixel size of ∼0.72, 0.83, or 0.93 Å/pixel. Movie frame alignment was conducted using MotionCor2 with dose weighting ^52^.

### 3D#reconstruction of the AAV9:Fab complexes

For the initial reconstruction using icosahedral averaging cisTEM was utilized as reported previously ^53,54^. Briefly, the motion-corrected micrographs were imported into the program, and their contrast transfer functions (CTFs) calculated. Micrographs of poor quality were removed. The capsid-Fab complexes were automatically picked using a characteristic particle radius of 135 Å and the individual capsid images extracted. These were then sorted via 2D classification into 50 classes. Classes containing impurities were discarded. The Ab-Initio 3D function was utilized to generate an initial map using 10% of the particles. This map was further refined using the automatic refinement function with default settings. The resolutions of the reported maps were determined at a Fourier shell correlation (FSC) criterion threshold of 0.143. The final electron density maps with imposed icosahedral averaging were sharpened using the pre-cut off B-factor value of −90 Å^2^ and variable post-cut off B-factor values of 0, 20, and 50 Å^2^.

In order to conduct localized reconstructions, the stack of individual particle images and the parameter file were exported in cisTEM to the Relion format. These particles were imported in Scipion with the pwem protocol and all further steps conducted in this program. As the first step the subparticles were defined by specifying the vector length from the center of the particle either along the 2-fold, 3-fold, or 5-fold symmetry axis (protocol: localrec – define subparticles). In the subsequent step the subparticles are filtered, keeping only unique particle images and removing view angles that significantly overlap with capsid (protocol: localrec – define subparticles). The individual images of the remaining subparticles are extracted (protocol: localrec – extract subparticles) and used for the reconstruction of an initial, low-resolution map with a C1 symmetry operator (protocol: relion – reconstruct). This map was then used as an input volume for the 3D classification protocol asking for at least four selected classes. During this step C2, C3, or C5 symmetry relaxation is activated. The resulting maps of the individual classes were inspected and the map with the highest resolution features for the capsid-Fab complex as well as the particles contributing to “good” classes selected for further refinement using the relion – 3D auto-refine protocol. The final map was sharpened using the relion – post-processing protocol.

### Docking of the AAV9 capsid and Fab models and scaling of the cryo-EM map

The atomic coordinates of the 60-meric AAV9 capsid (PDB accession number 3UX1) were docked into the icosahedral-averaged cryo-reconstructed density maps using the ‘fit in map’ subroutine in UCSF-Chimera ^55^. In order to identify the true pixel size of the maps a series of correlation coefficient (CC) calculations with different voxel sizes were conducted. The pixel size with the highest CC was utilized to resize the reconstructed map with the e2proc3d.py subroutine in EMAN2 ^56^. Subsequently the map was normalized and converted to the ccp4 format using MAPMAN ^57^. Structure models of the heavy and light chains of the Fabs were conducted with RoseTTAFold using the primary amino acid sequences ^58^. The resulting PDB files were docked into the icosahedral-averaged or localized reconstructed ccp4 maps by rigid body rotations and translations and by using the ‘Fit-in-map’ function in Chimera.

### Model refinement

The AAV9 capsid and the Fab light and heavy chain coordinates were refined in Coot ^59^ by manual building and utilizing the real-space-refinement subroutine to adjust side- and main-chains into the ccp4 map. The complexes were further refined against the maps using the real-space-refine subroutine in Phenix with default settings, which also provided refinement statistics (Table S1) ^60^. For the graphical representations of the maps and models Chimera and PyMol were utilized ^55,61^. The contact interactions for the AAV9-Fab complexes were analyzed using the online server PDBePISA ^62^.

### Enzyme-Linked Immunosorbent Assay

Enzyme-linked immunosorbent assays (ELISA) were performed as previously described^24^. Briefly, sera were assayed for reactivity to AAV9 or hAEV6 by ELISA. AAV vector preparations (50 µL) at a concentration of 2.5 x 10^10^ capsids/mL (in coating buffer (carbonate-bicarbonate buffer, Sigma-Aldrich) were used to coat 96-well polystyrene Maxisorp ELISA plates (Nunc) overnight at 4°C. The plates were washed 3 times with PBS + 0.05% Tween-20 (Sigma Aldrich) and then received 100 µL per well of blocking buffer (PBS + 5% skim milk + 0.05% Tween-20). After incubation at room temperature (RT) for 2 hours the plates were washed 3 times with PBS and received 50 µL per well of sera (serially diluted in blocking buffer as indicated with duplicate wells for each dilution). The plates were incubated for 2 hours at RT and washed 3 times with PBS before receiving 50 µL per well of horse radish peroxidase (HRP)-conjugated anti-human Fc specific IgG (Cell Signaling Technologies (Cat. CST32935) diluted 1:6,000 in blocking buffer). Plates were incubated for 1 hour at RT and washed 4 times before receiving 75 µL per well of 3,3′,5,5′-Tetramethylbenzidine (TMB, Sigma-Aldrich). Plates were incubated in the dark for 30 minutes at RT before reactions were stopped using 75 µL per well of 1M H_2_SO_4_. The absorbance of each well was measured at 450 nm wavelength using a VersaMax microplate reader (Molecular Devices, LLC). Duplicate wells containing no AAV served as background controls for each sera dilution. The mean value for each sample dilution was calculated for wells both with (foreground) and without coated vector (background) and the endpoint titer was determined as the lowest dilution where this ratio is >2.0. The limit of sensitivity for each assay is indicated in the graphs.

## Data availability

The cryo-EM reconstructed density maps and models built for the AAV9-Fab complexes were deposited in the Electron Microscopy Data Bank (EMDB). The respective EMDB and PDB accession numbers are listed in table S1.

---

**Table S1:**
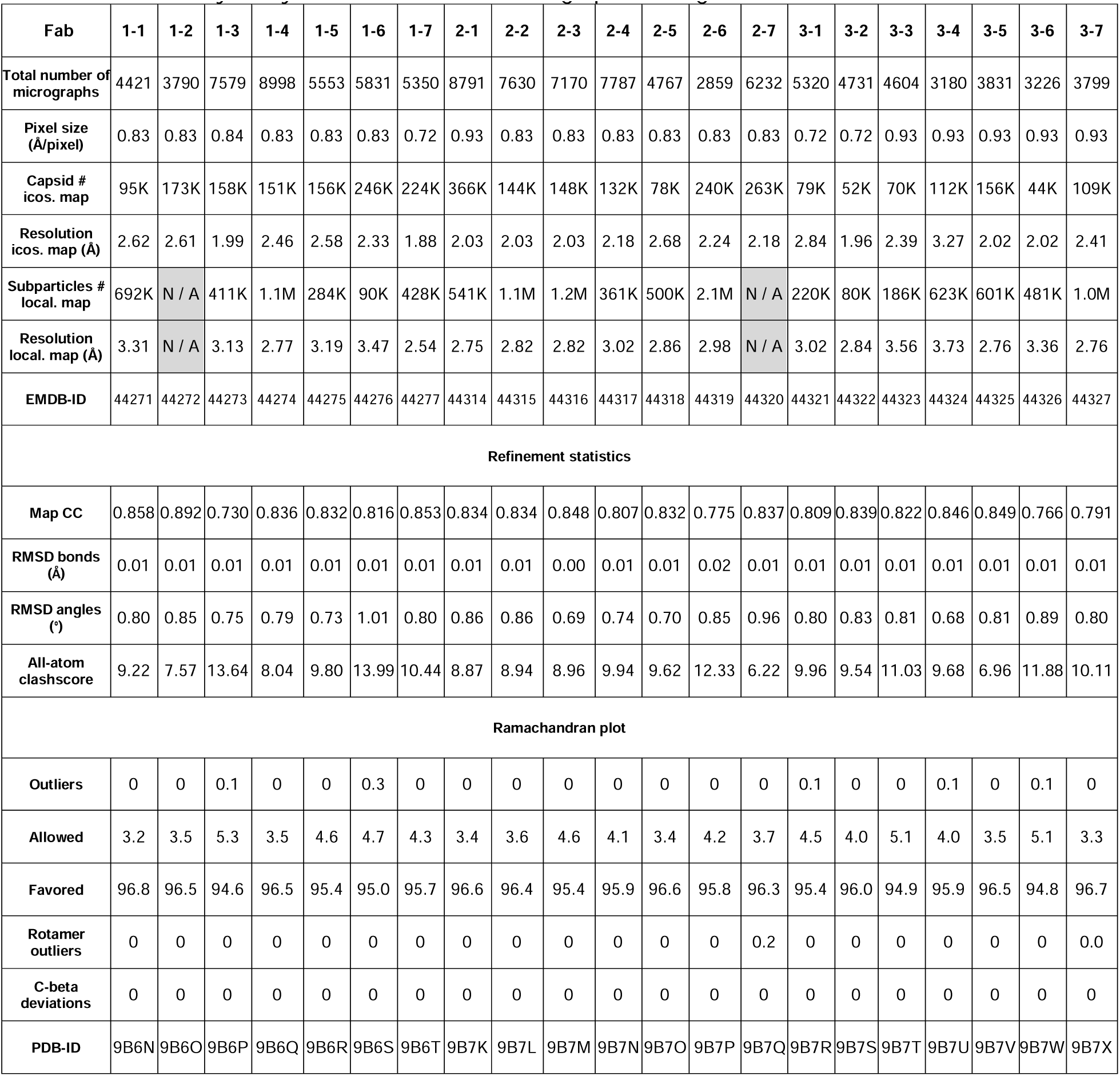
Summary of cryo-EM data collection, image processing, and refinement statistics.

**Table S2:**
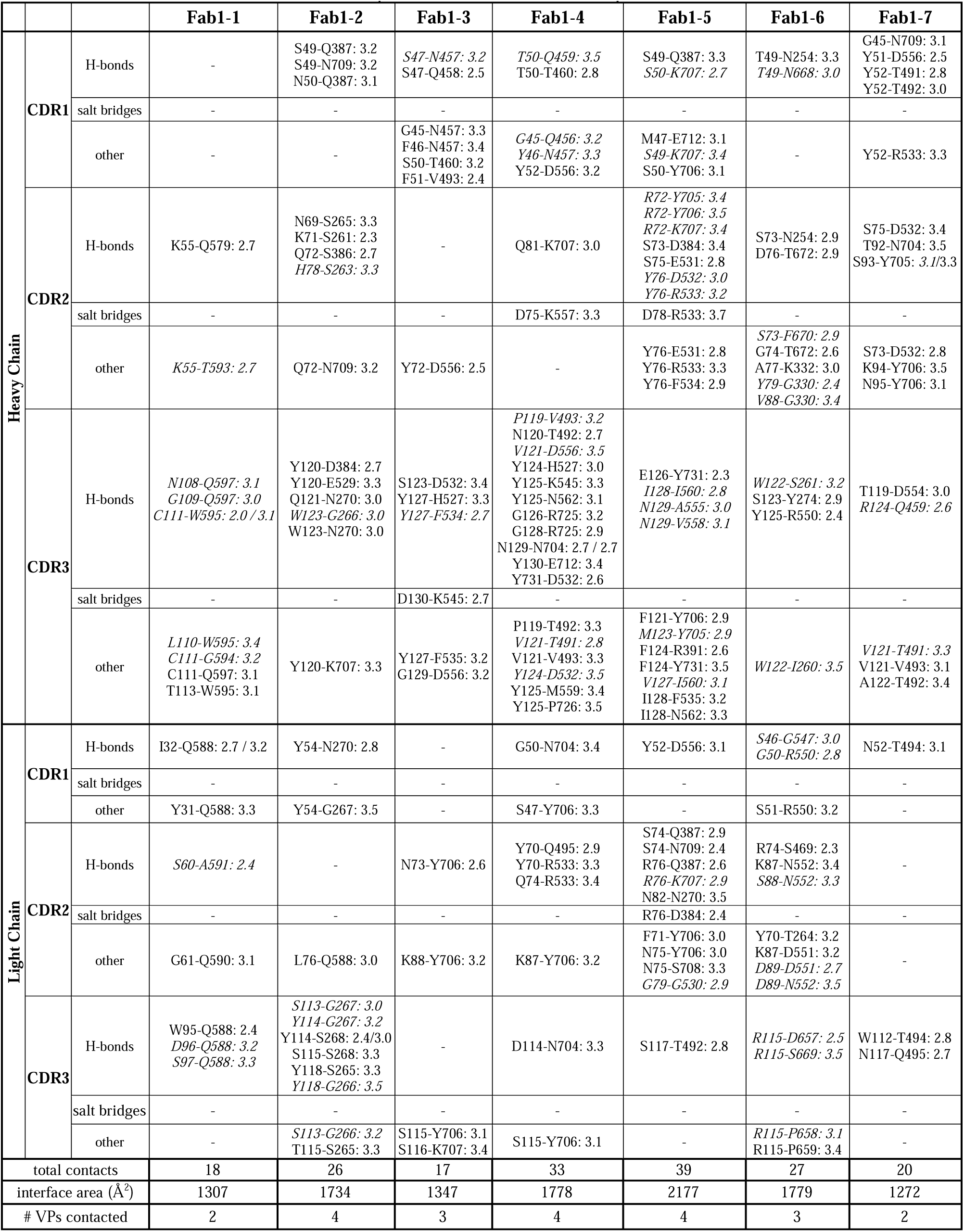
Contacts of the Fabs from patient 1 to the AAV9 capsid.

**Table S3:**
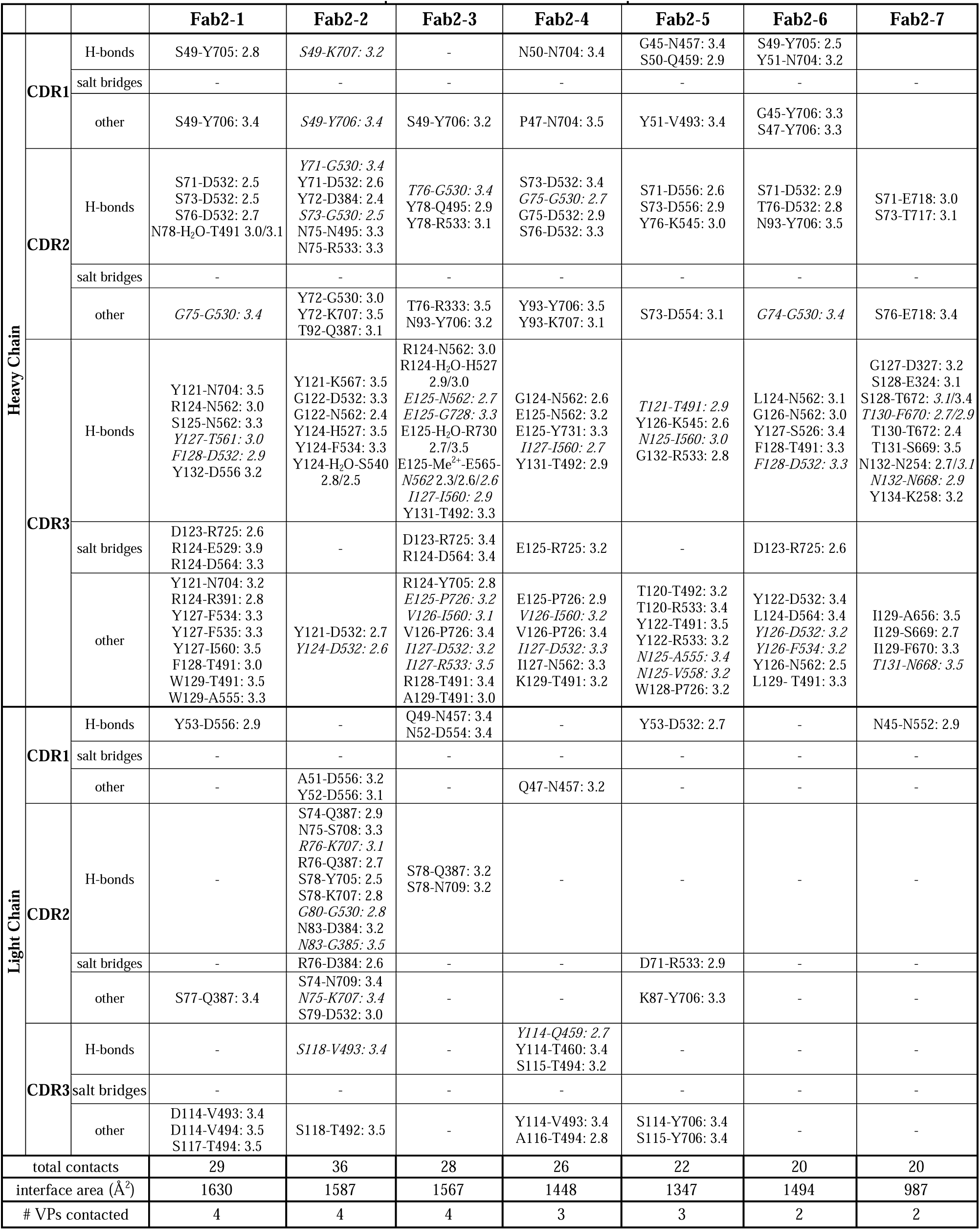
Contacts of the Fabs from patient 2 to the AAV9 capsid.

**Table S4:**
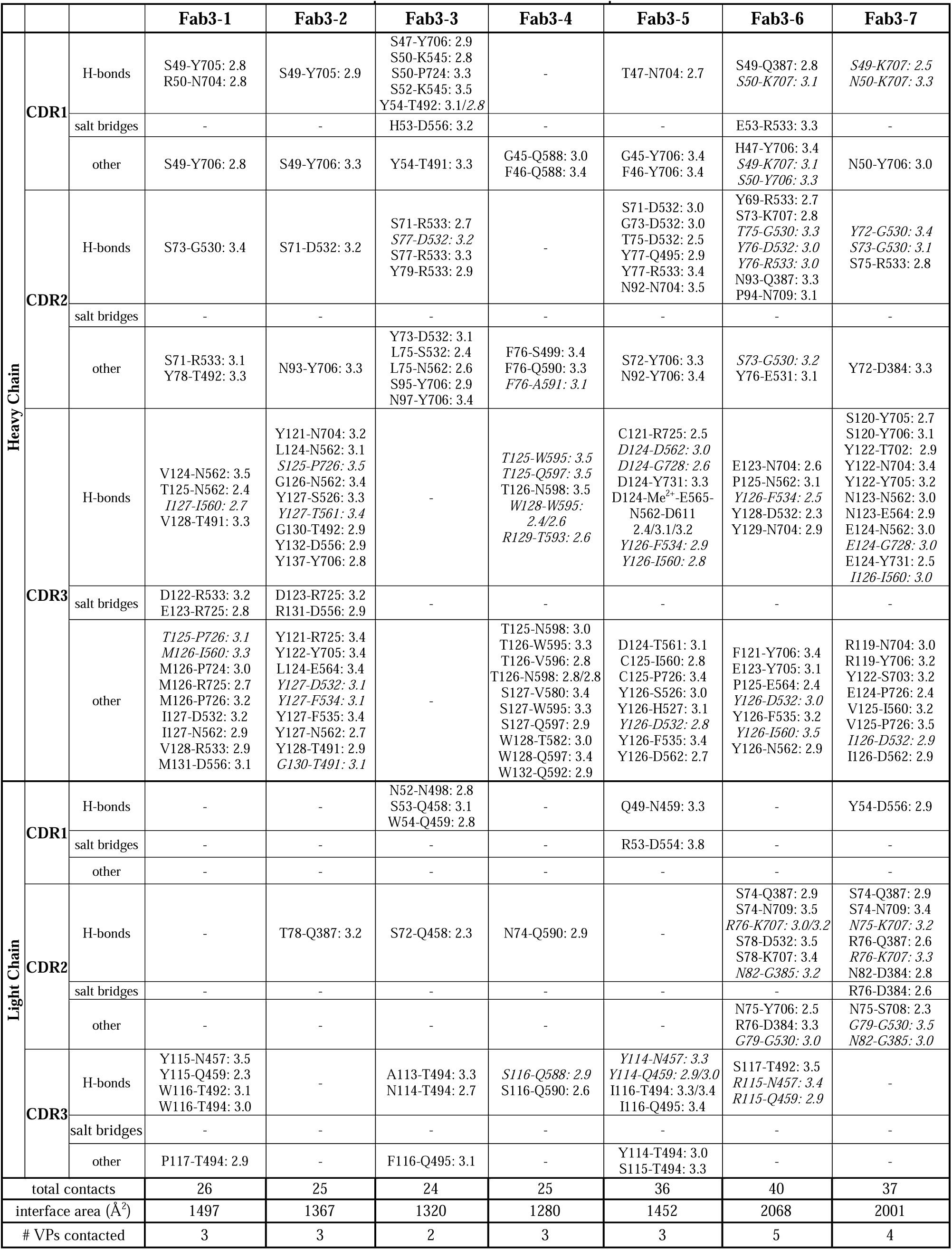
Contacts of the Fabs from patient 3 to the AAV9 capsid.

**Table S5:** Capsid variants tested against Fab1-2, Fab1-6, and Fab2-7.

| **Capsid variant** | **<u>Fab1-2</u>** | **<u>Fab1-6</u>** | **<u>Fab2-7</u>** |
| --- | --- | --- | --- |
| N254H | not tested | does not escape | does not escape |
| K258R | does not escape | does not escape | does not escape |
| S261A | ---- Capsid variant not infectious ---- |  |  |
| N262T | does not escape | not tested | does not escape |
| T264A | ---- Capsid variant not infectious ---- |  |  |
| G266S | ---- Capsid variant not infectious ---- |  |  |
| G267S | ---- Capsid variant not infectious ---- |  |  |
| Ins-A267 | ---- Capsid variant not infectious ---- |  |  |
| N270Q | ---- Capsid variant not infectious ---- |  |  |
| D384Q | ---- Capsid variant not infectious ---- |  |  |
| S386R | ---- Capsid variant not infectious ---- |  |  |
| D441N | does not escape | does not escape | does not escape |
| A502N | ---- Capsid variant not infectious ---- |  |  |
| G530E | does not escape | not tested | does not escape |
| G530N/D532N | does not escape | not tested | does not escape |
| R533S | does not escape | not tested | does not escape |
| G549Q | does not escape | does not escape | does not escape |
| R550S | ---- Capsid variant not infectious ---- |  |  |
| N552A/V553A | does not escape | does not escape | does not escape |
| I560F | does not escape | not tested | does not escape |
| A656Q | ---- Capsid variant not infectious ---- |  |  |
| A656G/P658L | not tested | does not escape | not tested |
| S669Q | not tested | does not escape | not tested |
| N704Q | ---- Capsid variant not infectious ---- |  |  |
| Y706D/N709I | does not escape | does not escape | does not escape |

## Figure legends

**Figure S1.** Phylogenetic relationship of the 2-fold binding Fabs. **A)** A dendrogram of the Fab V_H_ chains was generated at https://ngphylogeny.fr/ using their primary amino acid sequence. The groups of the 2-fold Fabs are color-coded (orange = group A, green = B, blue = C, beige =D). B) The lengths of the CDR3_H_ are plotted for each group. C) Dendrogram as in (A) for the Fab V_L_ chains.

**Figure S2.** Bivalent cation in the binding interface. A bivalent cation, e.g. calcium, was identified in the binding interface of Fab2-3 (and Fab3-5) stabilized by multiple residues and two ordered water molecules. Left: The atomic model is shown inside the cryo-EM density map at a σ- threshold of 2. Right: The distances of the interactions in the model are provided.

**Figure S3.** The human mAbs only detect native capsids. Dot blots on AAV9 capsids loaded in their native condition and after incubation for 5 minutes at 95°C.

**Figure S4.** Contact frequency of the Fabs to the AAV9 capsid. AAV9 capsid surface representation with the binding frequency for the 3-fold (left), 5-fold (center), and 2/5-fold wall (right) binding Fabs are displayed which are colored according to the bar below.

**Figure S5.** Fab induced conformational changes at the 2-fold region. Atomic models of amino acid 704-707 adopt the same conformation in both symmetry-related VR-IX loops in absence of any Fab. In contrast, numerous variations of side-chain orientations for the residue range are observed following Fab binding.

**Figure S6.** AAV9-Fab contacts utilized for capsid engineering. **A)** Modeled AAV9-Fab1-1 interaction in VR-VIII, CDR1_H_, and CDR3_L_ inside the cryo-EM map. Substitution of Q588 to arginine (pink) disrupts the interaction between capsid and antibody. **B)** Modeled AAV9-Fab2-4 interaction in VR-IX, and CDR1_H_ inside the cryo-EM map. Substitution of Y706 to aspartic acid (pink) disrupts hydrophobic environment of the contact region. **C)** Modeled AAV9-Fab3-4 interaction in VR-VIII, and CDR3_H_ inside the cryo-EM map. Substitution of T582 to glutamine (pink) displaces the tryptophan on the Fab side and prevents Fab3-4 binding. **D)** Modeled AAV9-Fab3-2 interaction in VR-VII, and CDR3_H_ inside the cryo-EM map. Substitution of N562 to tyrosine (pink) clashes with CDR3_H_ and prevents Fab3-2 binding.

**Figure S7.** AAV9 capsid surface. Surface representations of the AAV9 capsid are shown colored according to the scale bars below. **A)** The capsid is colored based on its hydrophobicity. **B)** The capsid is colored based on its electrostatic potential. **C)** Capsid surface residues alanine, valine, leucine, and cysteine are colored red and tryptophan, tyrosine, methionine, and histidine are colored blue.

## Notes

### Competing Interest Statement

The authors have declared no competing interest.

