## Supplementary figures and images for "Structural characterization of antibody-responses from Zolgensma treatment provides the blueprint for the engineering of an AAV capsid suitable for redosing"

### Supplemental Figure 1

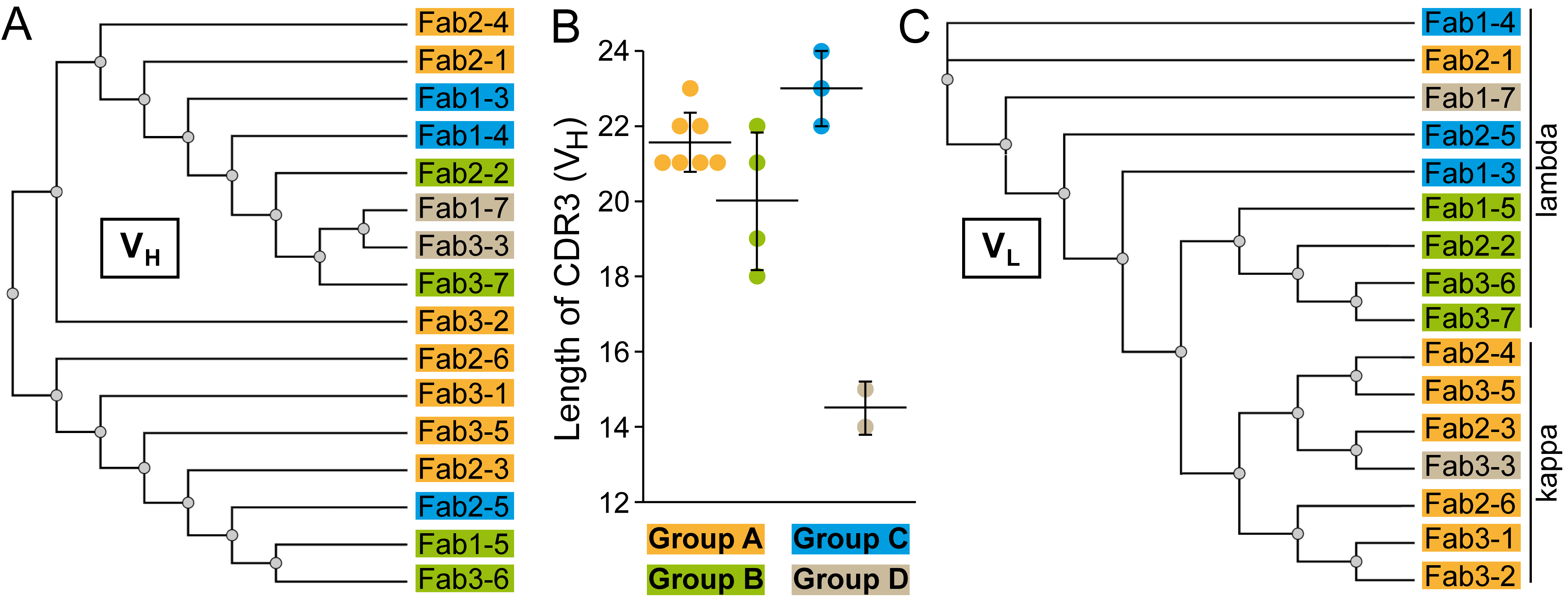

### Supplemental Figure 2

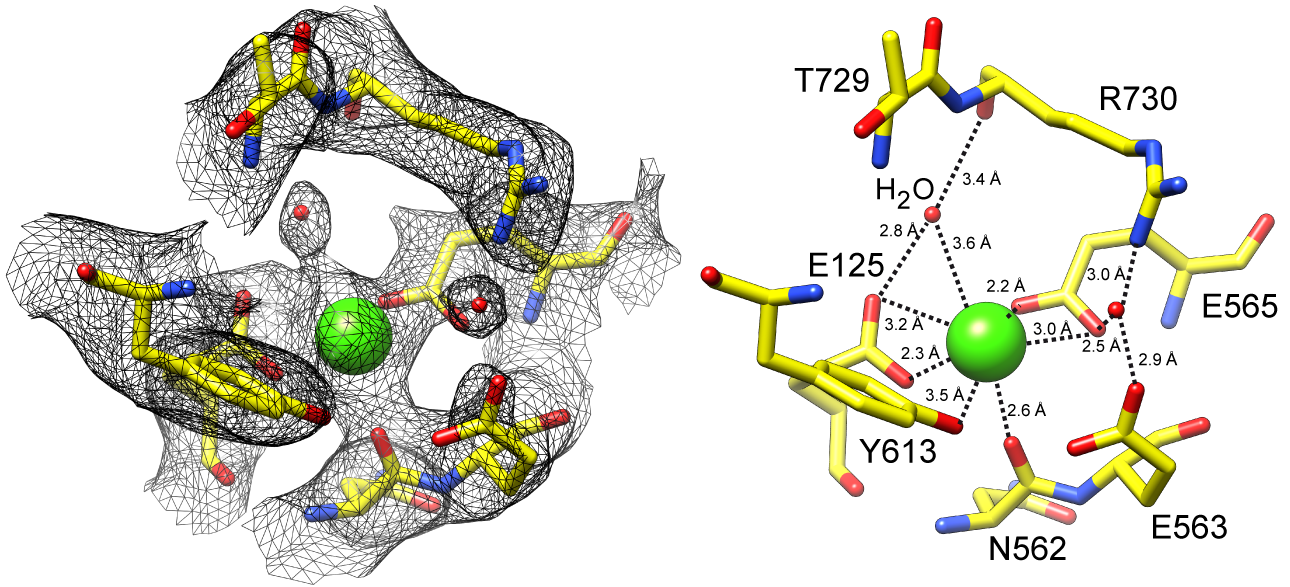

### Supplemental Figure 3

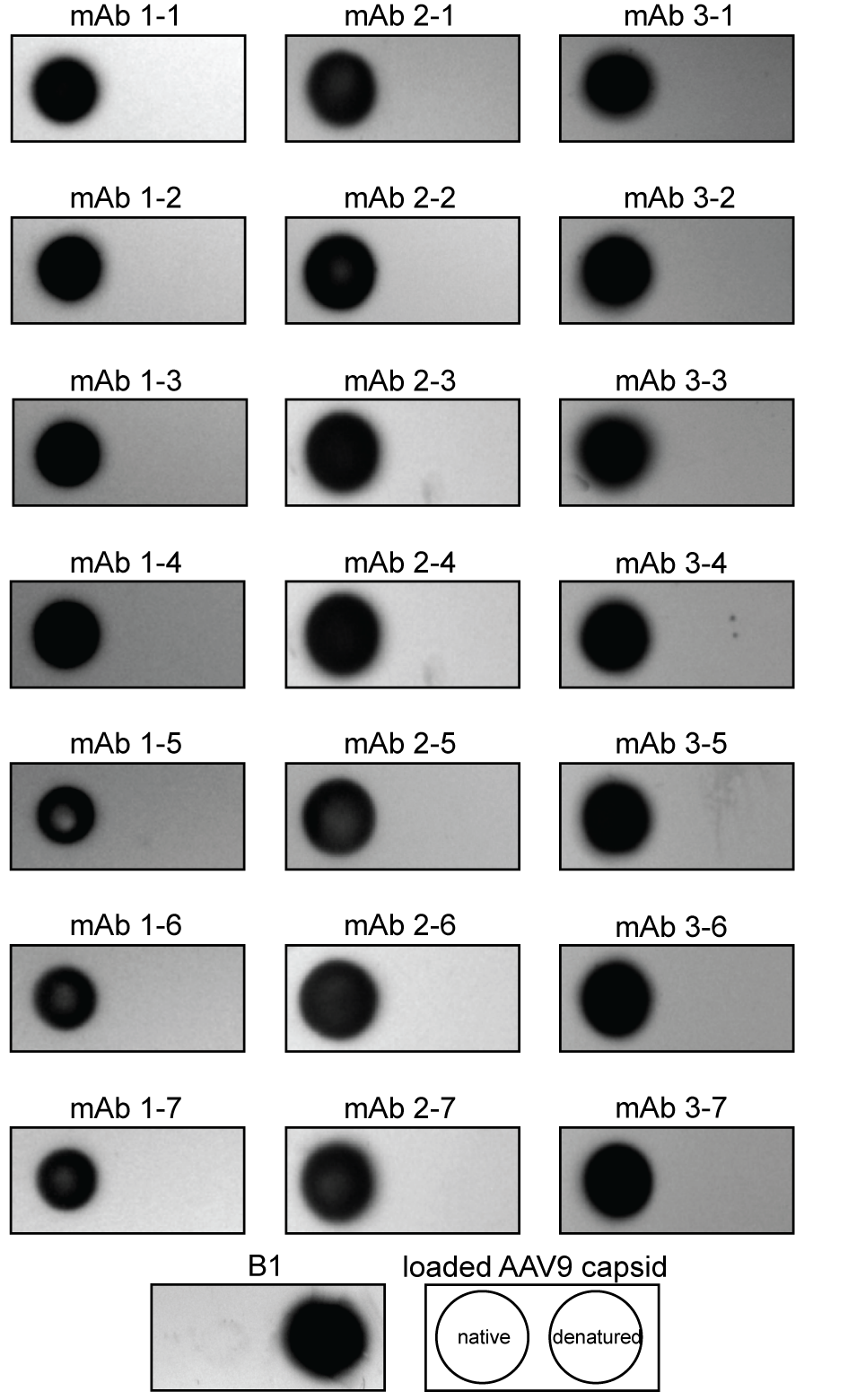

### Supplemental Figure 4

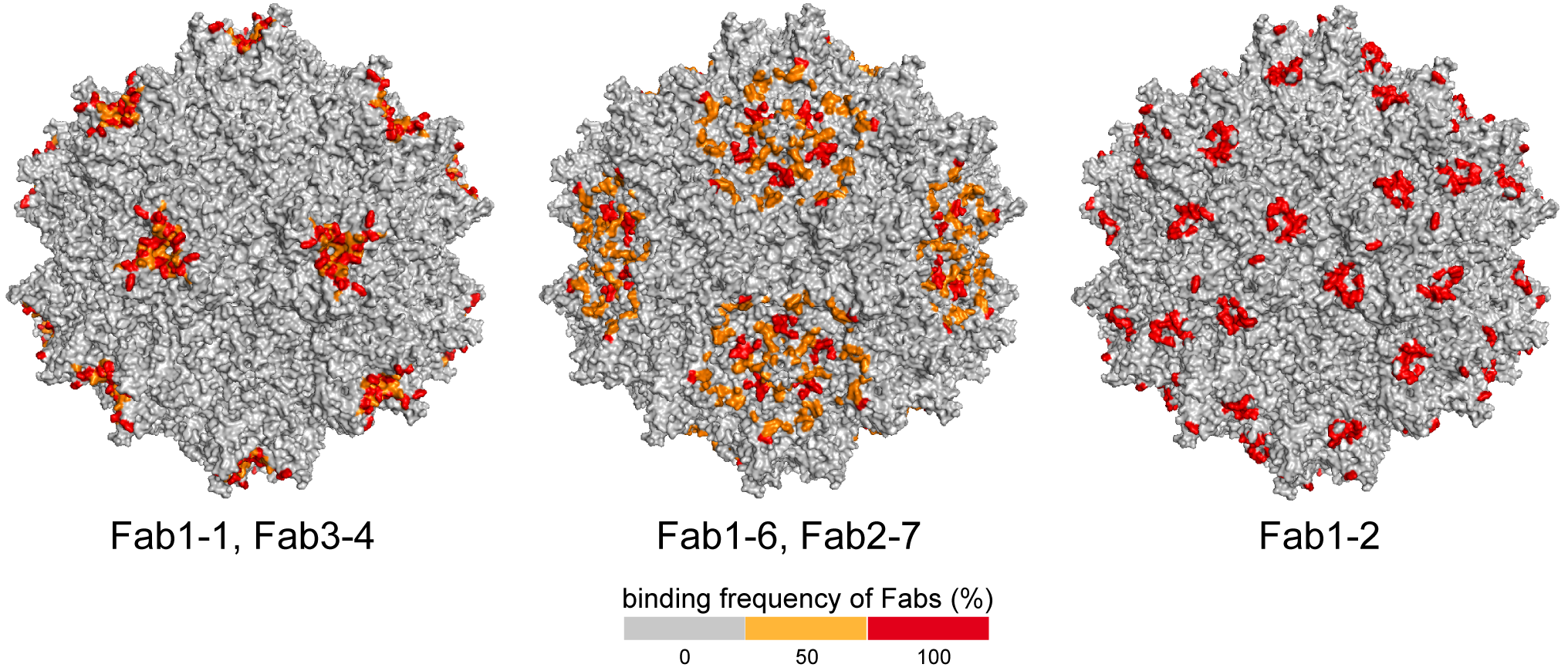

### Supplemental Figure 5

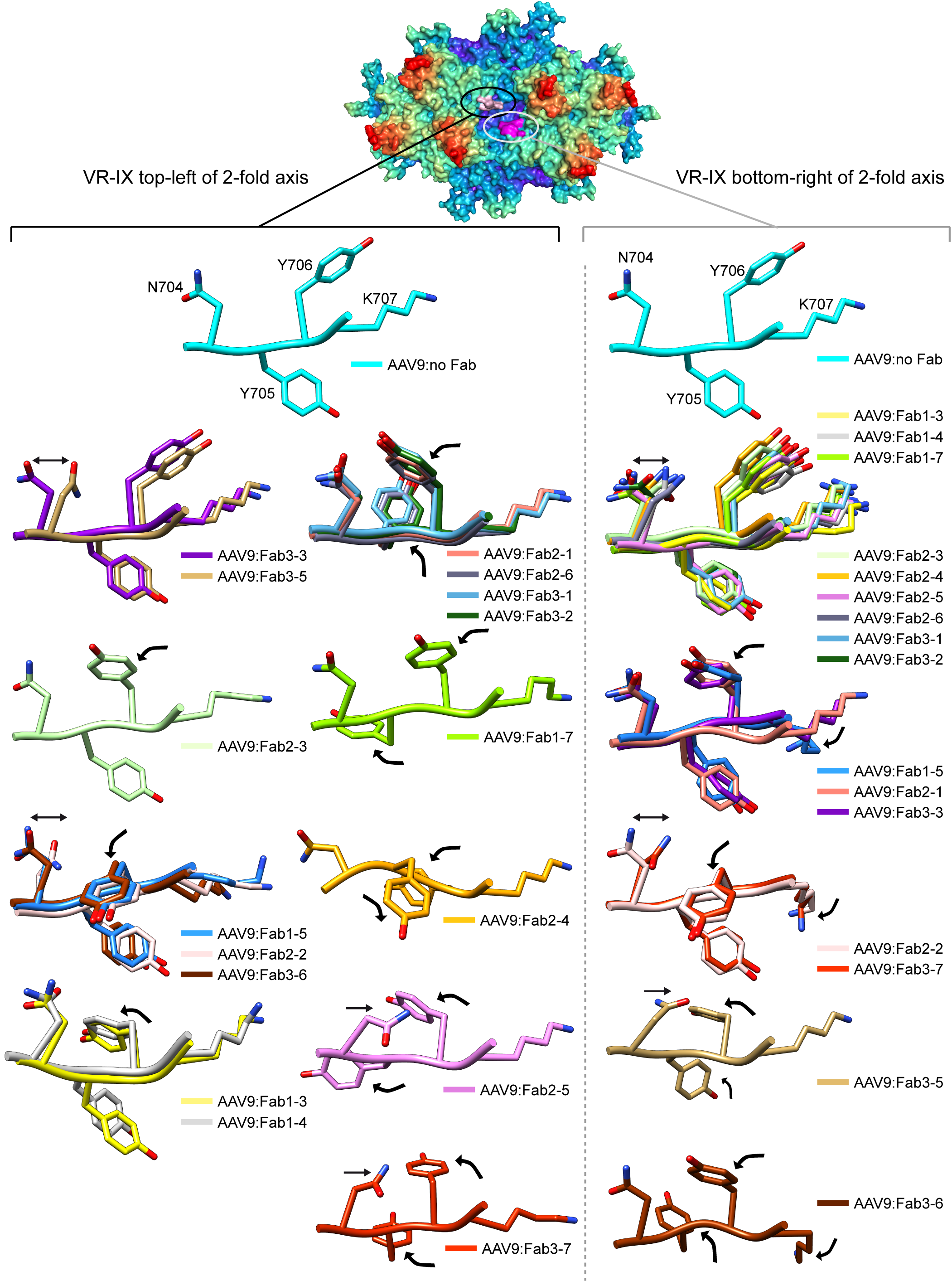

### Supplemental Figure 6

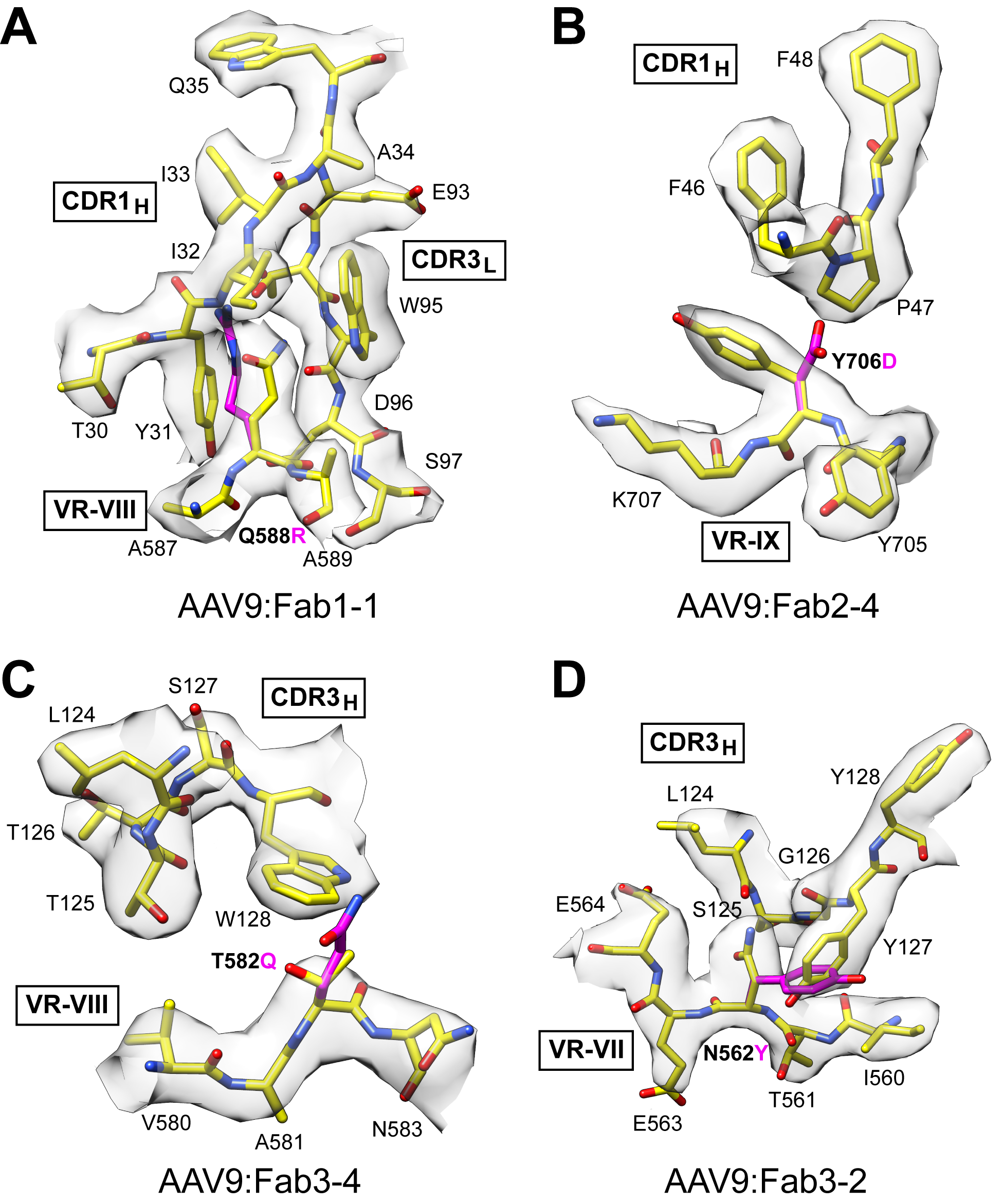

### Supplemental Figure 7

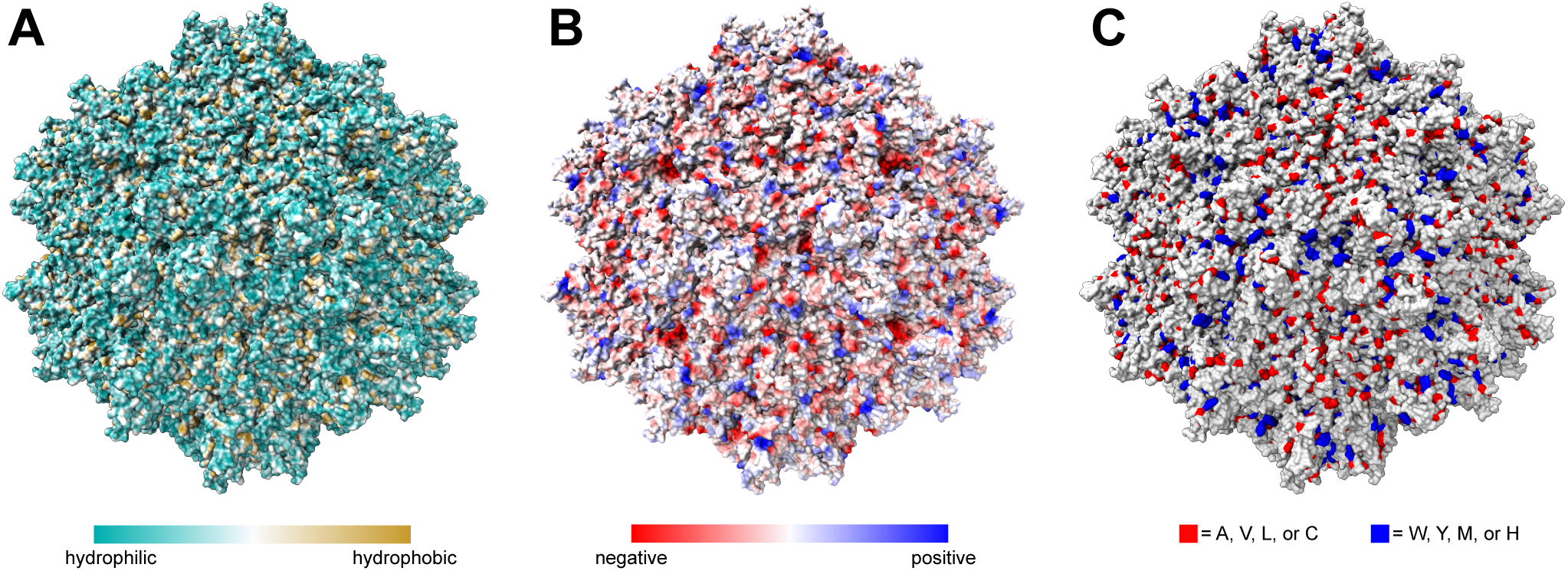
